# EnteroBase: Hierarchical clustering of 100,000s of bacterial genomes into species/sub-species and populations

**DOI:** 10.1101/2022.01.11.475882

**Authors:** Mark Achtman, Zhemin Zhou, Jane Charlesworth, Laura Baxter

**Author notes:** Pasteurien College, Soochow University, No.199 Ren’ai Road, Suzhou, Jiangsu 215123, China (Zhemin Zhou). Corresponding authors: MA,; ZZ.

## Abstract

The definition of bacterial species is traditionally a taxonomic issue while bacterial populations are identified by population genetics. These assignments are species specific, and depend on the practitioner. Legacy multilocus sequence typing is commonly used to identify sequence types (STs) and clusters (ST Complexes). However, these approaches are not adequate for the millions of genomic sequences from bacterial pathogens that have been generated since 2012. EnteroBase (http://enterobase.warwick.ac.uk) automatically clusters core genome MLST allelic profiles into hierarchical clusters (HierCC) after assembling annotated draft genomes from short read sequences. HierCC clusters span core sequence diversity from the species level down to individual transmission chains. Here we evaluate HierCC’s ability to correctly assign 100,000s of genomes to the species/subspecies and population levels for *Salmonella, Escherichia, Clostridoides, Yersinia, Vibrio* and *Streptococcus*. HierCC assignments were more consistent with maximum-likelihood super-trees of core SNPs or presence/absence of accessory genes than classical taxonomic assignments or 95% ANI. However, neither HierCC nor ANI were uniformly consistent with classical taxonomy of *Streptococcus.* HierCC was also consistent with legacy eBGs/ST Complexes in *Salmonella* or *Escherichia* and with O serogroups in *Salmonella*. Thus, EnteroBase HierCC supports the automated identification of and assignment to species/subspecies and populations for multiple genera.

## 1. Introduction

> “Microbiologists need large databases to identify and communicate about clusters of related bacteria, … Such databases should contain the reconstructed genomes of bacterial isolates,… together with metadata describing their sources and phenotypic properties… and we are currently developing EnteroBase, a genome-based successor to the *E. coli* and *S. enterica* MLST databases.” Excerpted from Achtman and Zhou, 2014 [1].

The Linnean system of genus and species designations was applied to bacterial nomenclature in the late 19^th^ century, soon after their ability to cause infectious diseases had been recognised. These taxonomic labels initially reflected disease specificity and phenotypic similarities. Over the following decades, phenotypic criteria for distinguishing bacterial taxa were extended to include serologically distinct groupings and even differential sensitivity to bacteriophages. A primary goal for taxonomic designations for bacterial pathogens was that they be useful for epidemiology and clinical diagnoses. As a result, serovars of *Salmonella enterica*, which differ by dominant epitopes on lipopolysaccharide and flagella, were each assigned a distinct species designation, often referring to the disease syndrome and host e.g. *Salmonella typhimurium* [2]. Similarly, even though *Yersinia pestis* is a clone of *Yersinia pseudotuberculosis* [3, 4], it was designated as a distinct species because *Y. pestis* causes plague whereas *Y. pseudotuberculosis* causes gastroenteritis. The use of species designations for *Salmonella* serovars is now disparaged, although it is still in common use [5], but *Y. pestis* has retained its species designation.

Percentage DNA-DNA hybridization levels became the “Gold Standard” metric for new taxonomic designations after 1987 [6]. DNA-DNA hybridization reflects genomic relationships but distinctive phenotypic differences have remained a requirement for the definition of a novel species until the present. Indeed, the international committee which controls taxonomic designations continues to reject the validity of taxonomic designations based solely on DNA similarities [7, 8].

### a) ANI and MLST

An alternative genomic approach for taxonomic designations was proposed in 2005 by Konstantinidis [9], namely the definition of species on the basis of average nucleotide identity (ANI). Pairs of genomes with 95% ANI usually belong to the same species whereas pairs with lower ANI values belong to different species. Computational methods based on k-mer searches, such as FastANI [10], can rapidly perform ANI-like calculations on large numbers of genomes, and these calculations are now routinely used by bioinformaticians for taxonomic assignments. An approach based on ANI is enticing because it could provide defined criteria for the definition of a species across all Bacteria, and allow species assignment based exclusively on genome sequences. However, the use of ANI for species assignments has been criticized because it does not completely correlate with DNA-DNA hybridization [11, 12]. Furthermore, multiple taxa that are designated as single species each contain multiple 95%ANI groups [13, 14]. For example, strains of *Streptococcus mitis* colonise the human oropharyngeal tract, have similar phenotypes and are considered to represent a single species. However, *S. mitis* encompasses myriad, genetically distinct strains [15–17], and encompasses at least 44 distinct 95% ANI clusters [17]. Other problematic genera include *Pseudomonas* [12, 18] and *Aeromonas* [19]. Furthermore, large databases including 100,000s of genomes per genus would struggle to implement methods such as pairwise clustering by ANI because each new entry would require testing against all genomes. There is also no consensus on ANI criteria for recognizing lower taxonomic entities, such as sub-species and populations. Indeed, there is not even a consensus on the definitive properties of what constitutes a bacterial species [20].

Bottom-up population genetic approaches such as multilocus sequence typing (MLST) can provide an alternative to top-down taxonomy, and deal efficiently with large numbers of bacterial strains. Legacy MLST based on the sequence differences of several housekeeping genes was introduced in 1998 [21], and has now been applied to more than 100 bacterial species [22]. MLST defines sequence types (STs), consisting of unique integer designations for each unique sequence (allele) of each of the MLST loci. Some STs mark individual clones with special pathogenic properties and which seem to have arisen fairly recently, such as *Escherichia coli* ST131 which is a globally prominent cause of urinary tract infection (UTI) and invasive disease [23]. Similarly, *Salmonella enterica* subsp. *enterica* serovar Typhimurium ST313 is a common cause of extra-intestinal, invasive salmonellosis in Africa [24]. Higher order clusters of related STs are also well known. Such clusters can be recognized by eBurst analyses [25], and are referred to as ST Complexes in *E. coli* [26] and eBGs (eBurst groups) in *S. enterica* [27]. ST Complexes and eBGs seem to reflect natural populations, but their broad properties are still not well understood.

The principles of legacy MLST were extended to rMLST, which uses sequences of 53 universal bacterial genes encoding ribosomal proteins [28] and provides a universal MLST scheme for all Bacteria [28]. For both *Escherichia* and *Salmonella,* rMLST offers a slight improvement in resolution over legacy MLST, and the identification of *Salmonella* eBGs is reasonably consistent across both approaches [29]. rMLST has been used to identify tractable sets of representative genomes from large collections for calculating phylogenetic trees [28] and pan-genomes [17].

MLST has also been extended to cgMLST, which encompasses all the genes in a soft core genome [29–31], and cgMLST nomenclatures have been implemented for multiple bacterial genera [28]. The large number of loci encompassed by cgMLST schemes results in enormous numbers of cgMLST STs (cgSTs), but these can be clustered into groups of bacterial genomes at multiple levels of genomic diversity (hierarchical clustering) [32]. cgMLST provides considerably higher resolution than legacy MLST or rMLST, and initial analyses indicated that it is ideal for investigating transmission chains within single source outbreaks or for identifying population structures up to the genus level [28]. Here we focus on the automated assignments of genome assemblies from six genera of important bacterial pathogens to taxonomic and population structures by hierarchical clustering of cgSTs with the EnteroBase HierCC pipeline (Table 1) [32]. For five of those genera, HierCC is a full solution for taxonomic designations of species.

**Table 1.** Legacy data, newly assembled genomes and hierarchical cgST clusters in EnteroBase (09/2021). NOTE: # Genomic Loci: numbers of CDS whose alleles are automatically called for the wgMLST and cgMLST schemes. cut-off: maximum numbers of allelic differences within minimal spanning trees that define HC clusters of cgSTs. EnteroBase automatically calculates existing Legacy MLST STs according to legacy schemes for *Escherichia/Shigella* (Wirth *et al.*, 2006 [26], *Salmonella* (Achtman *et al.,* 2012 [27], and *Clostridioides* (Griffiths *et al.,* 2010 [114]), but does assign new STs nor does it maintain a database of legacy data from ABI sequencing. Additional public databases are presented by EnteroBase for *Helicobacter* (>3500 genomes) and *Moraxella* (>2350 genomes), but these lack a cgMLST scheme. EnteroBase also has a database for >80,000 *Mycobacterium* genomes, but this currently (March, 2022) lacks a cgMLST scheme and a curator, and is not yet publicly available.

| Genus | Legacy MLST | cgMLST | # Genomic Loci |  | # HierCC clusters<br>(HierCC level) |  |  |
| --- | --- | --- | --- | --- | --- | --- | --- |
|  | (# strains) | (# genomes) | wgMLST | cgMLST | ST<br>Complexes | Lineages | Species/ Sub-species |
| *Salmonella* | 4,930 | 312,196 | 21,065 | 3,002 | 3648 (900) | 2185<br>(2000) | 15 (2850) |
| *Escherichia/*<br>*Shigella* | 9,525 | 175,256 | 25,002 | 2,513 | 1379 (1100) | 343<br>(2000) | 16 (2350) |
| *Streptococcus* |  | 76,718 | 33,887 | 372 | 3681 (100) |  | 132 (363) |
| *Clostridioides* |  | 23,197 | 11,490 | 2,556 | 299 (950) |  | 12 (2500) |
| *Vibrio* |  | 12,267 | 152,249 | 1,128 | 2000 (800) |  | 155 (1090) |
| *Yersinia* | 4,286 | 4,023 | 19,591 | 1,553 | 451 (600) |  | 34 (1490) |

### b) EnteroBase and HierCC

EnteroBase (http://enterobase.warwick.ac.uk) includes tools for automatic downloading of short read sequences and their metadata from the public domain, assembly into draft genomes and population genetic analyses of the core and accessory genomes (Table 2) [28]. Draft genomes are annotated according to genus-wide pan-genome schemes created with PEPPAN [17]. In September 2021, EnteroBase contained over 600,000 draft genomes from the six genera in Table 1, and likely provides a nearly comprehensive overview of their global diversity. Many of these samples reflect a focus on food-borne disease in the U.S. and United Kingdom, but this bias is increasingly being reduced by the global sources of many genomes (Supplemental Text, Sample Bias).

**Table 2.** EnteroBase-related software tools that support the analysis of large numbers of bacterial genomes. NOTE: PEPPAN is a stand-alone program that was used to generate the wgMLST schemes used in EnteroBase. SPARSE is a second stand-alone program that has been used to extract taxon-specific reads from metagenomes of ancient DNA that were then used with EToKi to define pseudo-MAGs that were uploaded to EnteroBase for phylogenetic comparisons. HierCC was used as a stand-alone program in development mode to define an initial set of clusters from a representative set of genomes. Subsequent assignments to existing clusters or to novel clusters were performed automatically within EnteroBase using pHierCCin production mode. BlastFrost is used by EnteroBase to determine the presence/absence of toxins and other pathovar characteristics of gastrointestinal pathogens within *E. coli*, and to assign gastrointestinal pathovar designations.

| Name | Purpose | Citation | URL for Stand-alone version |
| --- | --- | --- | --- |
| GrapeTree | GUI for depicting and analysing minimal spanning and NJ trees of character data | [115] | https://github.com/achtman-lab/GrapeTree |
| PEPPAN | Pan-genome calculation including pseudo-genes from numerous representatives of an entire genus | [17] | https://github.com/zheminzhou/PEPPAN |
| SPARSE | Assignment of metagenomic reads to individual taxa | [116,117] | https://github.com/zheminzhou/SPARSE |
| EToKi | EnteroBase toolkit of useful functions and pipelines | [28] | https://github.com/zheminzhou/EToKi |
| BlastFrost | Efficient k-mer based search for DNA sequences in large genomic datasets | [118] | https://github.com/nluhmann/BlastFrost |
| HierCC | Automated hierarchical clustering of cgSTs to existing or novel clusters | [32] | https://github.com/zheminzhou/pHierCC |

One of the primary goals for EnteroBase was a hierarchical overview of the population structure of the genera in Table 1. We therefore developed HierCC [32] based on cgMLST assignments to support the rapid recognition and detailed investigation of differing levels of population structure. EnteroBase reports cluster assignments and designations at 10-13 levels of allelic differences for all six genera [28] (Table 1). HC5 – HC10 clusters with maximal internal pair-wise distances within minimal spanning trees of 5 or 10 alleles, respectively, have been used to identify short-term, single source outbreaks of *S. enterica* and *E. coli/Shigella* that extended to multiple European countries [33–37].. Pathogen species can also include higher level clusters, which can correspond to somewhat more distantly related bacterial populations that cause endemic or epidemic disease over longer time periods in one or more countries [38–41]. EnteroBase HierCC has even been used to classify all *Shigella* [42], which correspond to discrete Lineages of *E. coli* [26, 43], and to define novel sub-species of *S. enterica* [44].

Classical taxonomic approaches for assigning individual strains and genomes to species depend on human expertise, and are not suitable for automated pipelines in real-time databases such as EnteroBase. This problem is acute because many short read sequences in ENA have incorrect taxonomic assignments, or none at all, and phenotypic distinctions are inappropriate for databases containing 100,000s of genomes. Zhou *et al.* compared taxonomic designations with peak normalized mutual information and silhouette scores for HierCC clusters, and identified HierCC levels that corresponded with well-defined species/sub-species in each genus [32]. For example, for *Salmonella* with a total of 3002 loci in the cgMLST scheme, HC2850 (94.9%) was chosen as the optimal HC level for identifying species and sub-species. The optimal HC levels for the other five genera with cgMLST schemes ranged from HC363 – HC2500 (93.6-97.8% of all cgMLST alleles) (Table 1). Here we compare the consistency of those HierCC assignments with assignments based on classical taxonomic approaches and 95% ANI. The results illustrate problems with classical approaches and with 95% ANI, and we conclude that HierCC is preferable for automated assignment of genomes to species/subspecies for those genera. However, neither ANI nor HierCC is universally satisfactory for species/sub-species assignments within *Streptococcus.* This manuscript also provides an initial overview of the abilities of HierCC to assign genomes to populations and Lineages [45, 46] within *Salmonella* and *Escherichia*/*Shigella*, and compares those assignments with the distributions of O antigens within lipopolysaccharide.

## Results

### a. Species and sub-species

In order to test the efficacy of HierCC at identifying species and sub-species, we extracted collections of representative genomes from all six genera in EnteroBase for which HierCC clusters had been implemented (Table 3). We wished to calculate maximum likelihood phylogenetic topologies of these genomes based on their core genome SNPs or the presence/absence of accessory genes from the pan-genome. Such ML trees are very slow to calculate with large datasets, but disjointed tree merging (DTM) within a divide-and-conquer approach allows large phylogenetic trees to be calculated in a reasonable time [47]. We therefore developed cgMLSA (see Methods in Supplemental Text), a novel DTM approach based on ASTRID [48] and ASTRAL [49] that enabled the calculations of ML super-trees from up to 10,000 representative genomes within hours to days. and applied it to each of the datasets. The resulting ML super-trees were annotated with taxonomic designations and cluster designations according to a 95% ANI cutoff calculated with FastANI [10] (95% ANI clusters). We then compared those annotations with cluster assignments based on the species-specific HierCC levels in Table 1.

**Table 3.** Parameters of datasets used for calculating ML super-trees. Each dataset consists of genomes representing the entire diversity of a genus as described in Methods (Supplemental Text).

| Genus | # Type strains | Total # Genomes | # Strict core genes | # Accessory genes | # Soft core genes (cgMLST) | # SNPs |
| --- | --- | --- | --- | --- | --- | --- |
| *Salmonella* | 7 | 10,002 | 1409 | 19,486 | 3002 | 1,410,331 |
| *Escherichia* (EcoRPlus) | 8 | 9479 | 134 | 20,294 | 2513 | 1,172,200 |
| *Escherichia* reps | 2 | 967 | 192 | 17,983 | 2513 | 971,516 |
| *Streptococcus* | 84 | 5937 | 90 | 31,630 | 372 | 263,080 |
| *Clostridioides* | 1 | 6725 | 515 | 10,975 | 2556 | 725,240 |
| *Vibrio* | 119 | 5032 | 124 | 146,041 | 1128 | 776,439 |
| *Yersinia* | 23 | 1847 | 744 | 18,799 | 1553 | 656,143 |

#### Salmonella

The topologies of ML super-trees based on 1,410,331 core SNPs from 10,002 representative genomes were in large part concordant with traditional taxonomic assignments, and with clustering according to 95% ANI (Fig. 1A) or HC2850 (Fig. 1B). The three sets of assignments were also largely concordant with the topology of an ML super-tree based on the presence or absence of accessory genes (https://enterobase.warwick.ac.uk/ms_tree/53258). HC2850 and 95%ANI distinguished *S. enterica* from *S. bongori*, and from former subspecies IIIa, which was recently designated *S. arizonae* by Pearce *et al*. [50]. HC2850 also identified another new *Salmonella* species, cluster HC2850_215890, and that identification was confirmed by 95% ANI. HC2850_215890 consists of five strains that have been isolated from humans since 2018 in the U.K., and a gene for gene comparison indicated that roughly half of their core genes were more similar to *S. enterica* and the other half to *S. bongori*.

**Figure 1.**
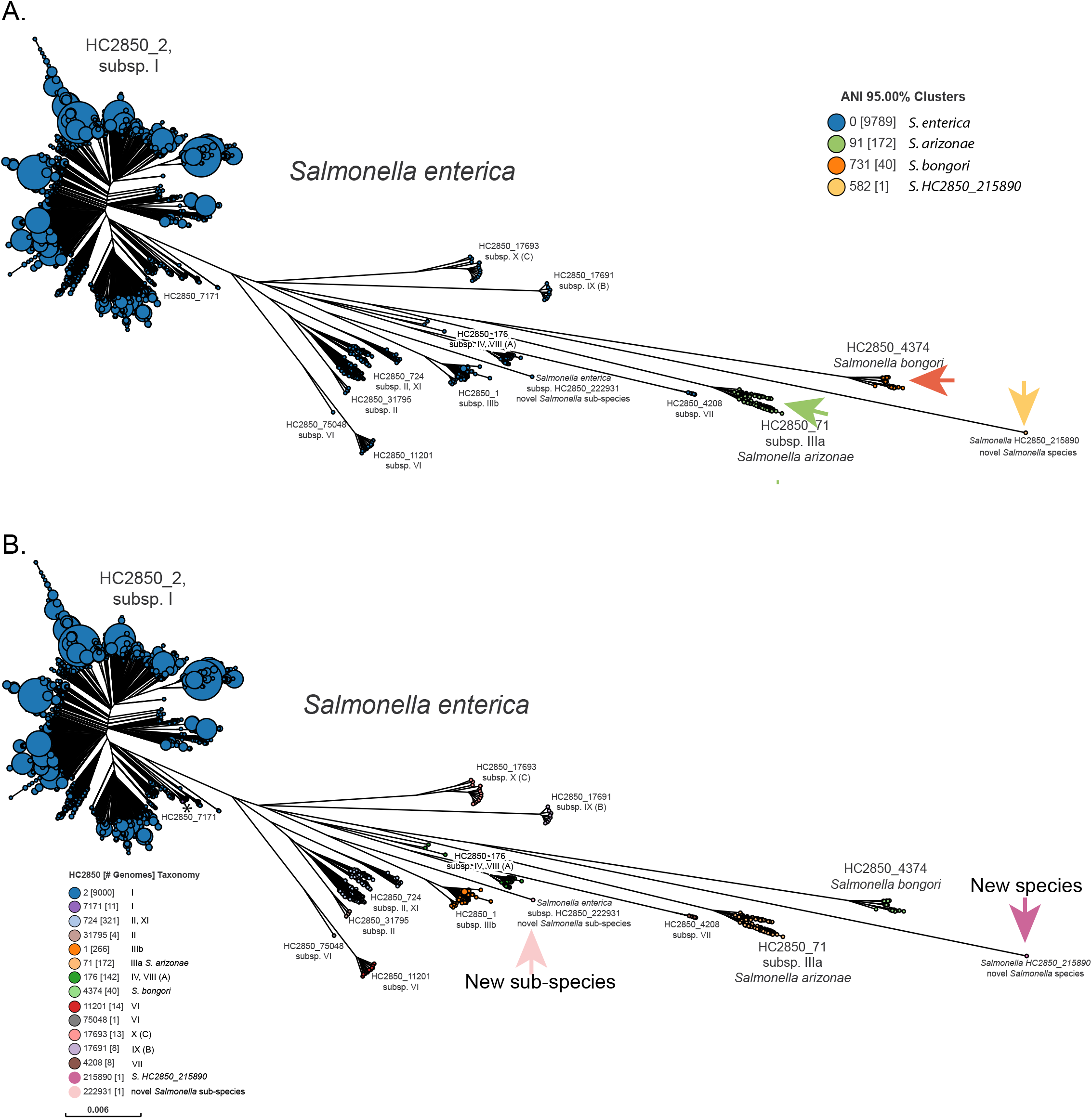
Species and sub-species assignments within *Salmonella* with HierCC and ANI. ML super-tree of 1,410,331 SNPs among 3002 core genes from 10,002 representative genomes of *Salmonella*. Former sub-species IIIa is designated *S. arizonae* in accordance with Pearce et al., 2021 [48]. A. Partitions differentiated by ANI 95% clusters (legend) correspond to species (*S. enterica, S. bongori, S. arizonae, S.* HC2850_215890), as indicated by arrows, and sub-species are not differentiated. B. Partitions coloured by HC2850 clusters (legend). Arrows indicate a new species, HC2850_215890 (five strains from the U.K., 2018-2020), and a new sub-species, HC2850_222931 (one strain from France, 2018). All other HC2850 clusters correspond to species (*S. bongori* and *S. arizonae*) or subspecies, except for HC2850_7171 (starred), which is subsp. *enterica* (I) according to the ML tree. An interactive version of this GrapeTree rendition can be found at https://enterobase.warwick.ac.uk/ms_tree?tree_id=53257. The corresponding presence/absencetree can be found at https://enterobase.warwick.ac.uk/ms_tree?tree_id=53258.

95% ANI did not detect any subspecies structure within *S. enterica* whereas HC2850 differentiated a total of 12 clusters that corresponded to distinct phylogenetic branches in the ML super-tree (Fig. 1B). Most of these have been designated as subspecies by traditional taxonomy [44, 50], except for a singleton genome in HC2850_222931, which likely represents a novel subspecies. Despite the correct identification of so many sub-species, HC2850 did not support Pearce’s definition of subspecies XI, and did not distinguish those genomes from subsp. II. Similarly, HC2850 did not differentiate Pearce’s subsp. VIII from subsp. IV. The prior definition of these subspecies was largely dependent on branch topologies within phylogenetic trees based on rMLST loci, and was also contradicted by population statistics [50].

HierCC also differentiated HC2850_7171, which consists of a short phylogenetic branch within subsp. I that would not have been assigned to a subspecies according to its ML topology. Genomic comparisons of multiple single genes from genomes within HC2850_7171 suggest that HC2850_7171 may be a hybrid between subsp. I and II because some gene sequences were most similar to the former and others were most similar to the latter. With this sole possible exception, HierCC seems to be suitable for the automated detection of new *Salmonella* species and subspecies, and for routinely assigning novel genomes to the appropriate taxa.

#### Escherichia

In addition to *Escherichia coli, Escherichia* includes the named species *albertii* [51–53], *fergusonii* [54, 55]*, marmotae* [56, 57] and *ruysiae* [58]. And despite their apparently distinct genus and species names, *Shigella boydii, Shigella dysenteriae, Shigella flexneri* and *Shigella sonnei,* all common causes of dysentery, corresponds to phylogenetic lineages within *E. coli* [43] rather than to discrete taxonomic units. Still other, unusual *Escherichia* strains from lake and ocean water are associated with long phylogenetic branches, and were designated as ‘cryptic clades’ I-VIII by population geneticists [59–65]. The branch leading to Clade I is simply a long phylogenetic branch within *E. coli* [59]. However, Clade V encompasses *E. marmotae* [56, 57] and *E*. *ruysiae* consists of the union of clades III and IV [58]. Initial analyses with the EcoRPlus collection of 9479 genomes [28] (Table 3) yielded results that were compatible with these interpretations. Almost all *E. coli* or *Shigella* genomes were within HierCC cluster HC2350_1, and genomes with other taxonomic or Clade designations belonged to other HC2350 clusters. However, soon after the definition of the EcoRPlus collection, additional genomes of *Escherichia* which were related to Clade II were described from inter-tidal marine and fresh water sediments near Hong Kong [61, 66]. Furthermore, due to its numerical predominance, *E. coli* overshadows other *Escherichia* species and sub-species within EcoRPlus. We therefore created *Escherichia* reps in Jan 2021, a novel set of 967 representative genomes which included one genome from each of the 160 most common HC1100 clusters in HC2350_1 and all 807 genomes from other HC2350 clusters which existed in EnteroBase at that time.

Fig. 2 shows that *E. fergusonii* and *E. albertii* form discrete 95% ANI clusters as do *E. marmotae* and other Clade V genomes. As previously reported [58], *E. ruysiae* consists of the distinct clusters of Clades III and IV. Three distinct 95% ANI clusters were found within Clades II, VI, and VIII while the Hong Kong genomes represented multiple, related phylogenetic branches, one of which has previously been designated Clade VII. All *E. coli* and *Shigella* genomes are in a common 95% ANI group, as is Clade I. Comparable clustering results were also found in an ML tree of presence/absence of accessory genes, albeit with different topology of the deepest branches resulting in *E. fergusonii* and Clade VIII being most closely related to *E. coli* (https://enterobase.warwick.ac.uk/ms_tree?tree_id=71125). These observations indicate that the taxonomy and population genetic designations for *Escherichia* are incomplete, and also partially inconsistent.

**Figure 2.**
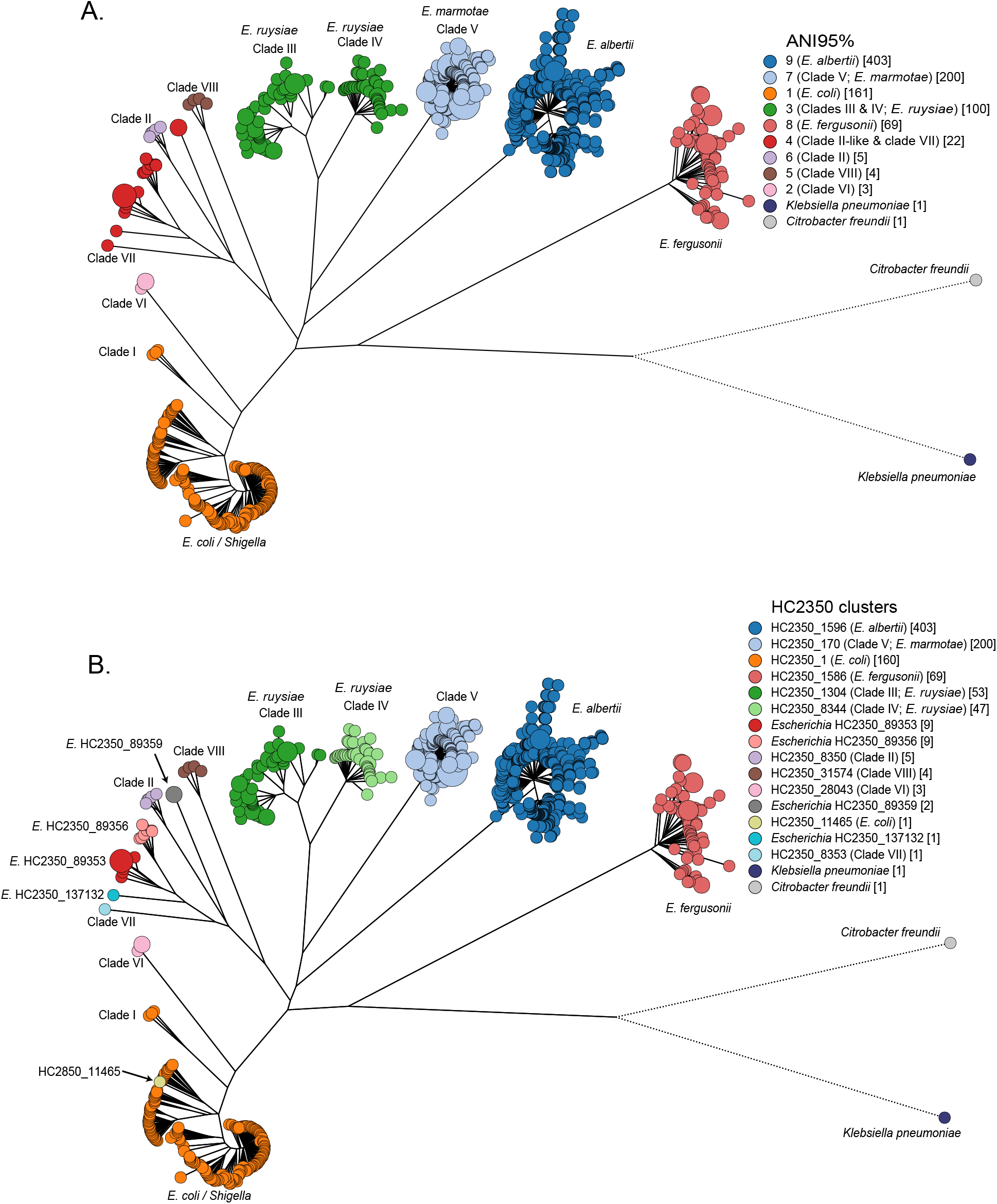
Maximum-likelihood core SNP tree of 967 *Escherichia* genomes consisting of one genome from each of 161 HC1100 clusters containing *E. coli* or *Shigella* as well as all 806 other *Escherichia* genomes in Enterobase as of November 2020. The tree is colored by A) pairwise FastANI values clustered at the 95% level and B) HC2350 cluster designations. The key legends indicates taxonomic designations in the literature which best match the cluster groupings. In part B, HC2350 cluster designations were used to mark novel taxonomic groupings in HC2350 clusters 89353, 89356, 89359 and 137132. An interactive version of this GrapeTree rendition can be found at https://enterobase.warwick.ac.uk/ms_tree?tree_id=52101. The corresponding presence/absence tree can be found at https://enterobase.warwick.ac.uk/ms_tree?tree_id=71125.

HierCC clustering of the same data provided a consistent and uniform nomenclature. Each discrete phylogenetic cluster received its own HC2850 designation, including Clades III and IV and four phylogenetic clusters among the Hong Kong isolates which were designated according to their HC cluster number: e.g. *Escherichia* HC2350_89353. *E. coli, Shigella* and Clade I all clustered together in HC2350_1, and the only discordance between topological clustering and HierCC clustering among the 967 genomes in *Escherichia* reps was a single genome which belonged to *E. coli* by phylogenetic topology but was assigned to HC2350_11465 by HierCC. These results were so convincing that HierCC groupings were used in late 2021 to curate and update all species designations for *Escherichia* entries within EnteroBase.

#### Clostridioides

*C. difficile* clustered distinctly from *C. mangenotii* according to 95% ANI, and ANI also distinguished five other clusters among genomes that were designated *C. difficile.* These clusters seem to be novel species according to their phylogenetic topology in the ML SNP super-tree (Fig. 3). HC2500 identified the same clusters, and also separated out two additional distinct clusters which resemble novel sub-species (arrows: HC2500_15334, HC2500_15408). A subset of the genomes in these clusters (Fig. 3) were assigned to cryptic clades C-I, CII and C-III by a recent publication [67], which also concluded that they represent novel species. Thus, HierCC is also suitable for the automated detection of new *Clostridioides* species and subspecies, and for routinely assigning novel genomes to the appropriate taxa.

**Figure 3.**
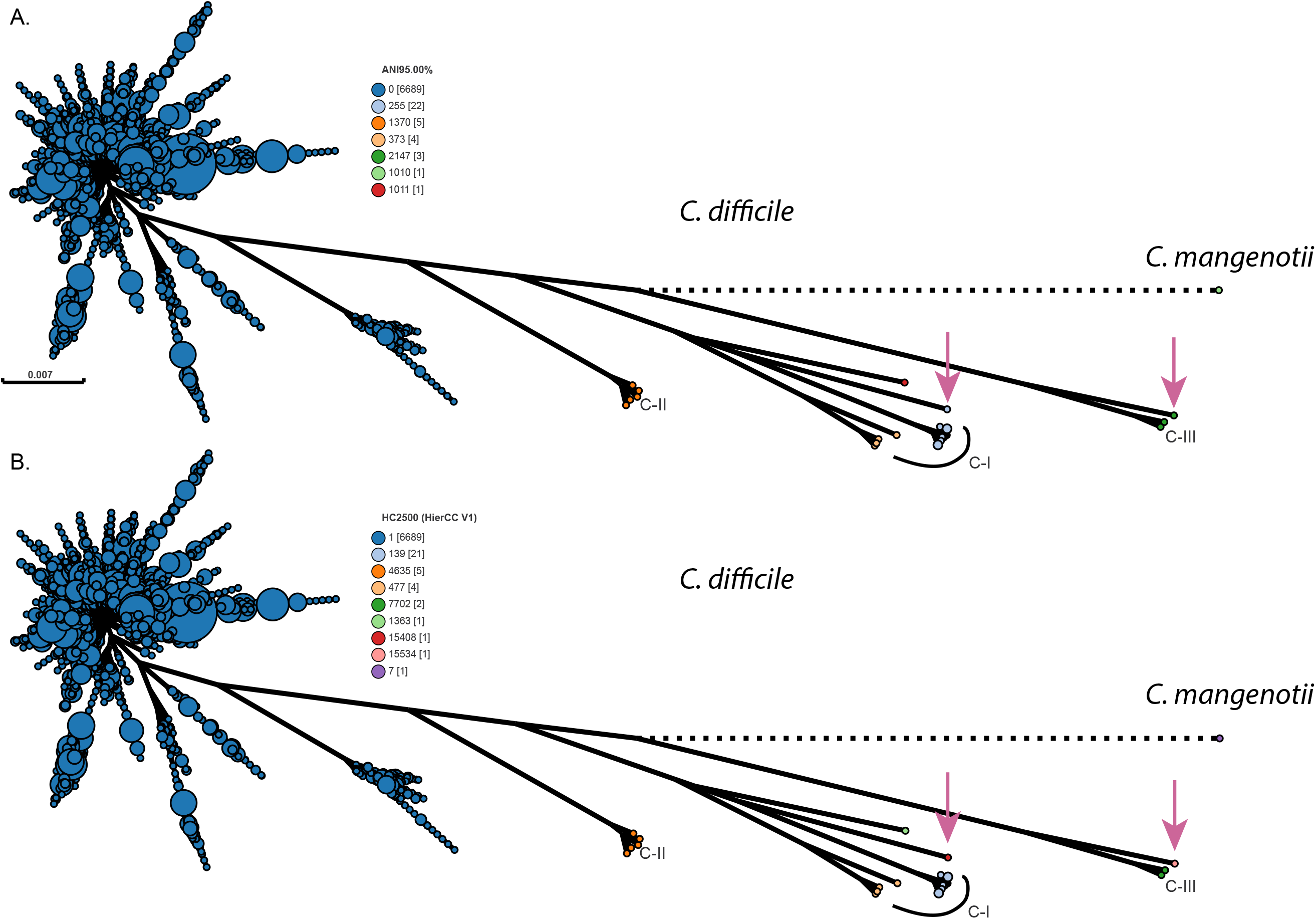
Species and sub-species assignments within *Clostridioides* according to ANI (A) and HierCC (B). ML super-tree of 725,240 SNPs among 2556 core genes from 6,724 representative genomes of *Clostridioides difficile* and one genome of *Clostridioides mangenotii*. A. ANI 95% clusters (legend) differentiate *C. mangenotii* (cluster 1010) and five other clusters (clusters 255, 1370, 373, 2147, 1011) from *C. difficile* (cluster 0). As indicated, four of these correspond to cryptic clades C-I [clusters 255, 373], C-II [cluster 1370] and C-III [cluster 2147]) according to Knight et al. [49]. Arrows indicate two additional clusters that were distinguished by HierCC in part B. B. Partitions coloured by HierCC assigns the same genomes as ANI to the corresponding HC2500 clusters, except that it defines HC2500_15334 and HC2500_15408 to one genome each, which might represent sub-species. An interactive version of this GrapeTree rendition can be found at https://enterobase.warwick.ac.uk/ms_tree?tree_id=53253 and a presence/absence tree can be found at https://enterobase.warwick.ac.uk/ms_tree?tree_id=53254.

#### Yersinia and Vibrio

Detailed results with *Yersinia* and *Vibrio* can be found in Supplemental Text. The general take-home lessons from those analyses resemble those summarized above: 95% ANI clustering is largely congruent with existing taxonomic designations, and HierCC identifies both species and sub-species with better resolution than 95% ANI. However, multiple, unanticipated taxonomic problems were evident in both genera. Firstly, ENA uses the strain designation as a species designation when a species is not specified during the submission process. The resulting public metadata is bloated with inaccurate species names. Secondly, multiple distinct phylogenetic branches were found without any species identifier. Thirdly, in multiple cases the species identifier did not correspond with ANI and/or HierCC, and comparisons with both the SNP and presence/absence ML topologies confirmed that the species designations in ENA were incorrect. We addressed these problems within EnteroBase by performing radical manual curation to the metadata to ensure that the taxonomic species designations correspond to the phylogenies and population structures. We also encountered multiple, problematical taxonomic designations which are described in the next paragraphs.

#### Yersinia

95% ANI supports the taxonomic convention that *Y. enterocolitica* represents a single species, and all genomes designated as *Y. enterocolitica* were in a single 95% ANI cluster. However, in accord with the conclusions by Reuter *et al.* [68], HierCC clustering assigned the non-pathogenic biotype 1A genomes to HC1490_73 and HC1490_764, the highly pathogenic biotype 1B to HC1490_2, and pathogenic biotypes 2-5 to HC1490_10. We therefore renamed these four groups by adding the HierCC cluster to the species name, e.g. *Y. enterocolitica* HC1490_2.

In contrast to *Y. enterocolitica,* the *Yersinia pseudotuberculosis* Complex corresponds to a single phylogenetic cluster according to both HC1490 and 95% ANI (Supplemental Fig. S1). Taxonomists have split these bacteria into *Y. pseudotuberculosis*, *Y. pestis, Y. similis* and *Y. wautersii* [69, 70]. DNA-DNA hybridization [71] and gene sequences of several housekeeping genes [4] previously demonstrated that *Y. pestis* is a clade of *Y. pseudotuberculosis,* and the new observations demonstrate that a distinct species status is not consistent with the genomic data for *Y. similis* and *Y. wautersii*. We have therefore downgraded all three taxa within EnteroBase to the category of subspecies of *Y. pseudotuberculosis,* and extended the designation of *Y. pseudotuberculosis* to *Y. pseudotuberculosis sensu stricto.* These assignments are not reflected by distinct HC1490 clusters, and HC1490 clustering can only be used to automatically assign new genomes to the *Y. pseudotuberculosis* Complex.

We also defined eight other novel species/sub-species designations within *Yersinia* by unique designations based on HierCC clusters: e.g. *Yersinia* HC1490_419. These clusters were previously unnamed, or had been incorrectly designated with the names of other species which formed distinct ANI and HC1490 clusters (SupplementalText, Fig. S1).

#### Vibrio

*Vibrio* encompassed 152 HC1090 clusters (Supplemental Fig. S2, SupplementalText), which corresponds to much greater taxonomic diversity than for the other genera dealt with above. The concordance between HC1090 and 95% ANI clusters was absolute for most clusters, including a large numbers of genomes from *V. cholerae, V. parahaemolyticus*, *V. vulnificans,* and *V. anguillarum* (Supplemental Table S1), and did not support the existence of additional sub-species. However, eleven HC1090 clusters each encompassed between two and five ANI clusters (Supplemental Table S2) and three ANI clusters each encompassed 2-3 HC1090 clusters (Supplemental Table S3). In order to support the automated assignment of genomes to named species, we implemented taxonomic assignments according to HC1090, and renamed the species of all genomes that were contradictory to this principle. The resulting dataset contains 109 HC1090 clusters with a unique, classical species designation as well as 43 other species level clusters designated as *Vibrio* HC1090_xxxx. Seventeen species names were eliminated because they were contradictory to the phylogenetic topologies or were incoherently applied (Supplemental Text, Supplemental Table 4). These taxonomic changes now permit future automated assignment of novel genomes to species designations, and the recognition of novel species, and have provided a clean and consistent basic taxonomy that can be progressively expanded.

Prior work has assigned many *Vibrio* species into so-called higher order clades of species on the basis of MLSA (multilocus sequence analysis) [72]. These clades are also apparent in the ML super-tree of core SNPs, and Fig. S2 indicates the three largest (Cholerae, Harveyii, Splendidus).

#### Streptococcus

For all five genera summarized above, clustering according to HierCC was largely concordant with the ML super-trees based on core SNPs or presence/absence of accessory genes. 95% ANI and taxonomic designations were also largely concordant with the phylogenetic trees, although to a lesser degree. Similar concordances with the trees based on core SNPs (Fig. 4) and presence/absence of accessory genes (Supplemental Fig. S3) were also found for a majority of the named species within *Streptococcus*. 102 HC363 clusters were concordant with 95% ANI clusters, and each of those HierCC clusters was specific for a single species after eliminating *S. milleri, S. periodonticum* and *S. ursoris* (Supplemental Table S5). These results also demonstrated strong agreement between the two methods with classical taxonomy. However, genomes from a large number of other species within *Streptococcus* were not clustered satisfactorily by either method.

**Figure 4.**
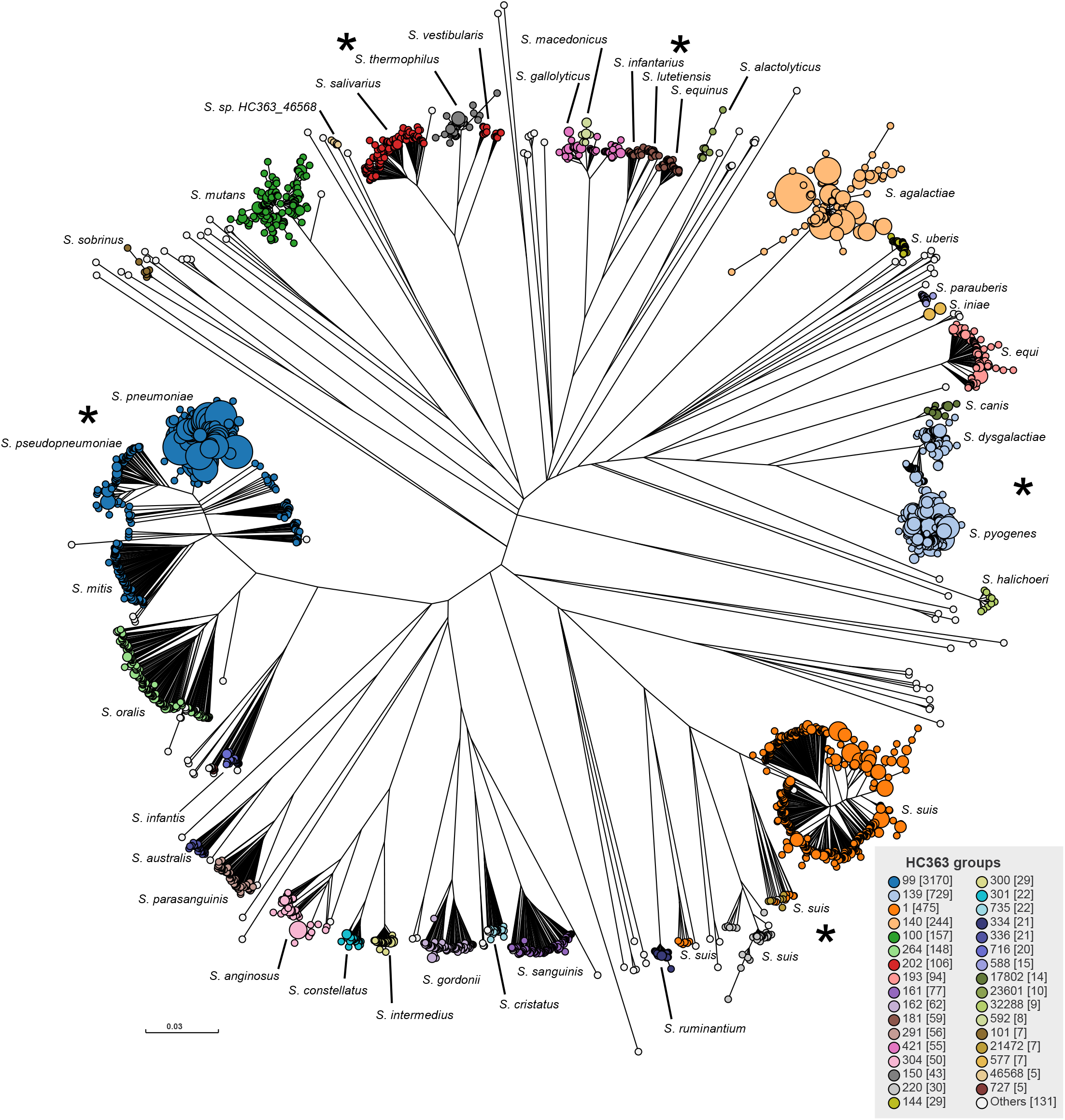
Comparison of HC363 clusters with taxonomic designations in *Streptococcus*. ML super-tree of 263,080 SNPs among 372 core genes from 5937 representative *Streptococcus* genomes. Species names are indicated next to the phylogenetic clusters where genomes mapped for type strains, and according to published metadata. Nodes are colored by HC363 clusters, and exceptional assignments are indicated by asterisks next to *S. pneumoniae* and *S. pseudopneumoniae,* which were both HC363_99; multiple phylogenetic and HierCC clusters grouped within *S. suis*; *S. salivarius* and *S. vestibularis*, which were both HC353_202; *S. lutetiensis and S. equinus*, which were both HC363_181; and *S. dysgalactiae and S. pyogenes*, which were both HC363_139. An interactive version of the GrapeTree rendition of the SNP tree can be found at https://enterobase.warwick.ac.uk/ms_tree?tree_id=53261 and the presence/absence tree at https://enterobase.warwick.ac.uk/ms_tree?tree_id=53262.

Earlier publications [15,16,73,74], had demonstrated that *Streptococcus mitis* and *oralis* each represents multiple related sequence clusters rather than two single species, and similar results were obtained with 95% ANI clustering [17]. These observations were confirmed by the current data. Of the 102 concordant HC363 and 95% ANI clusters, 23 clusters from *S. cristatus, S. infantis, S. mitis, S. oralis, S. sanguinis* or *S. suis* each corresponded to only a sub-set of the topological diversity within their species. Furthermore, single HC363 clusters from 18 species or groups of species were subdivided extensively by 95% ANI for a total of 155 discrepancies between the two methods (Supplemental Table S6). These discrepant clusters were within the species *S. suis, S. criceti, S. sanguinis*, *S. suis, S. oralis, S. hyointestinalis, S. parasanguinis, S. australis, S. infantis,* and *S. cristatus.* 95% ANI was not able to differentiate *S. pneumoniae* from *S. pseudopneumoniae*, *S. pyogenes* from *S. dysgalactiae, S. salivarius* from *S. vestibularis*, or between *S. equinus*, *S. infantarius,* and *S. lutentiensis.* Contrariwise, three other 95% ANI clusters each encompassed two HC363 clusters (Supplemental Table S7). HC363 could not distinguish *S. pneumoniae* or *S. pseudopneumoniae* from fifty-seven 95% ANI clusters within *S. mitis.* In addition to these problems, the eight additional species should not have been given a species name because their type strains belonged to one of these chaotic clusters. We conclude that numerous species have been defined on taxonomic grounds that cannot be correctly identified by either 95% ANI or HierCC, and that multiple discrepancies exist between the two approaches within *Streptococcus*.

### b. Populations, eBGs and Lineages

HierCC supports the identification of populations at multiple hierarchical levels ranging from the species/sub-species down to individual transmission chains. We also expected that intermediate HierCC cluster levels could reliably detect the natural populations defined by legacy MLST. Here we focus on examples of such populations from *Salmonella*, *E.coli/Shigella,* and *S. pneumonia,* taxa where they have been examined in greatest detail.

#### Salmonella

eBGs (eBurstGroups) defined by legacy MLST in *Salmonella* generally correspond to HierCC HC900 clusters [32]. In some cases, genetically related groups of eBGs correspond to HC2000 clusters [45]; we refer to those as Lineages. In contrast, none of the HierCC levels corresponded consistently with other, prior intra-species subdivisions of *S. enterica* subsp. *enterica* into lineages [75], Clades [76–79] or Branches [80]. This failure may reflect a high frequency of extensive homologous recombination at these deeper branches, which can obscure topological relationships [46, 80], whereas HC900 (and HC2000 clusters) diverged more recently, and have not yet undergone extensive recombination [46].

Traditional subgrouping nomenclature of *Salmonella* below the sub-species level predates DNA sequencing, and is predominantly based on serovar designations. Serovar designations consist of common names that have been assigned to a total of more than 2500 unique antigenic formulas based on epitopes within lipopolysaccharide (O antigen) and two alternately expressed flagellar subunits (H1, H2). These antigenic formulas are written in the form O epitopes:H1 epitopes:H2 epitopes in the revised Kauffmann-White scheme [81, 82], and O epitopes are summarized as O serogroups with distinct numeric designations, e.g. O:4 for the O group of serovar Typhimurium. Detailed metabolic maps for LPS synthesis have been elucidated for 47 O serogroups [83, 84]. Serogroups from natural isolates can be determined serologically by agglutination reactions with specific antisera or *in silico* from genomic sequences with the programs SeqSero [85] or SISTR [86]. In practice. each of these methods has an error rate of a few percent [27, 87].

Several hundred legacy eBGs within subsp. *enterica* were relatively uniform for serovar [27], with multiple exceptions, and many serovars were associated with multiple, apparently unrelated eBGs, likely due to the extensive exchange of genes encoding LPS and flagellar epitopes between eBGs early in their evolutionary history. We proposed that legacy MLST was a better metric for identifying the natural populations than serotyping, and could completely replace this traditional method [27]. Our new data indicate that HC900 clusters are even more reliable than legacy eBGs for recognizing natural populations within *Salmonella*.

Several phylogenetic comparisons indicated that HC900 clusters based on cgMLST are concordant with eBGs in *Salmonella* [32, 45]. Here we test this concordance quantitatively over a total of 319,490 genomes (Table 4). The concordance between eBG and HC900 clusters was 0.985 according to the Adjusted Mutual Information Score [AMI] (Table 4), a metric that is suitable for samples with a heterogeneous size distribution [88], and almost as high with the Adjusted Rand Index (ARI: 0.982), which is less suitable for heterogeneous data. Comparisons of eBG or HC900 with HC2000 clustering yielded slightly lower AMI scores (Table 4). We also tested whether HC900 clusters are uniform for serovar. In late 2019, we corrected false serological data in the metadata Serovar field in EnteroBase by manual curation of 790 HC900 clusters that contained at least 5 entries. 97% (770/790) of those HC900 clusters were uniform (≥95%) for serovar (Supplemental Table S8), as were most HC2000 clusters (437/473 clusters; 92%). The data indicated that the predominant O group was uniform over the multiple HC900 clusters within 95% (382/403) of HC2000 clusters within *S. enterica* subsp. *enterica* and 79% (58/70) of HC2000 clusters in other *Salmonella* species and sub-species (Fig. 5; Supplemental Table S9).

**Figure 5.**
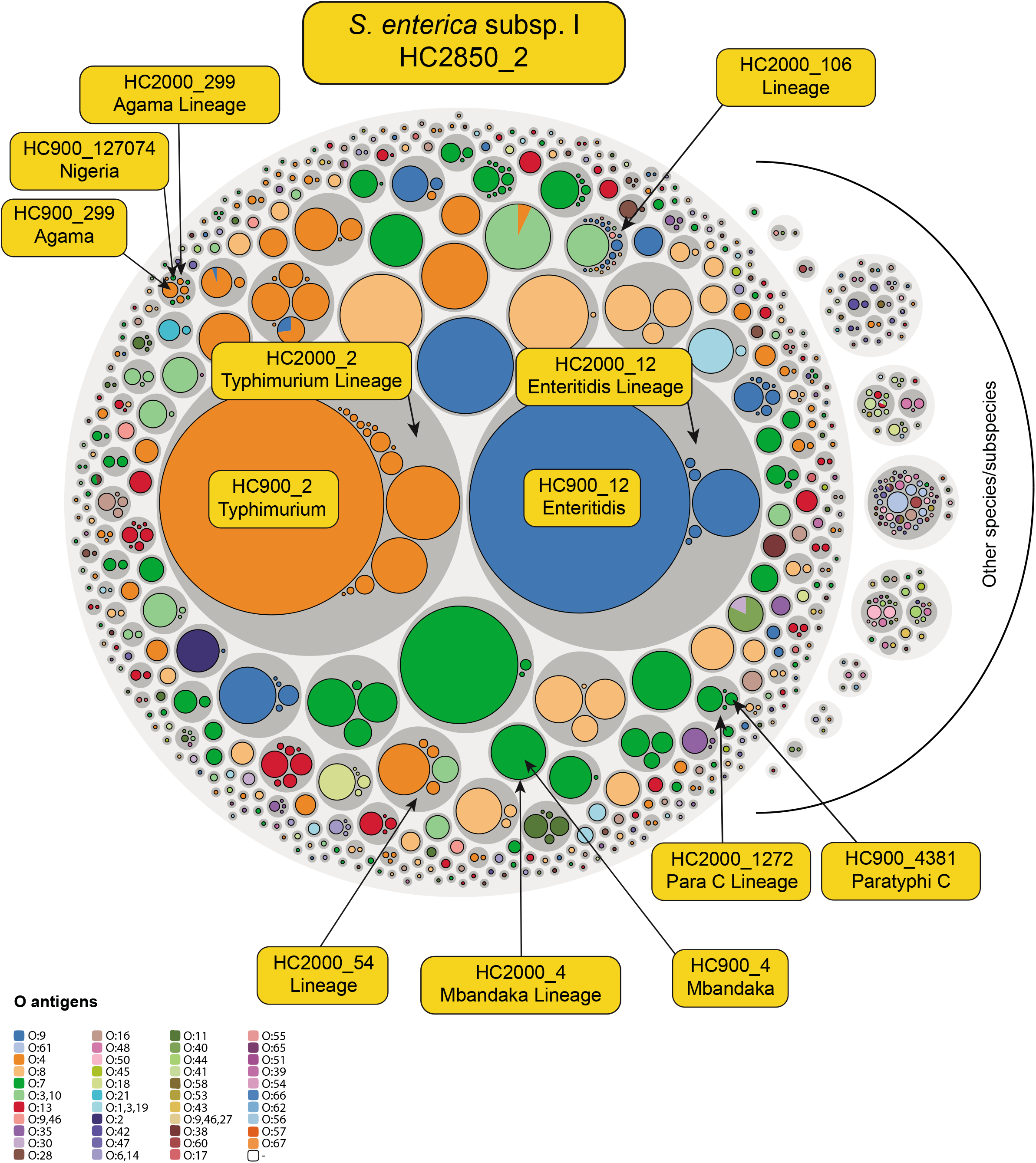
Hierarchical population structure of O serogroups in *Salmonella*. Hierarchical bubble plot of 310,901 *Salmonella* genomes in 790 HC900 clusters for which a consensus O serogroup could be deduced by metadata, or bioinformatic analyses with SeqSero V2 [85] or SISTR 1.1.1 [86]. Taxonomic level (HC level; colors): species/subspecies (HC2850; light grey circles), Lineages (HC2000; dark grey circles) and eBurst groups (HC900; O:group specific colors). Additional information is indicated by yellow text for selected HC2000 and HC900 circles which are specifically mentioned in the text. The diameters of HC900 circles are proportional to the numbers of genomes. An interactive version of this figure can be found at https://observablehq.com/@laurabaxter/salmonella-serovar-piechart from which the representation, raw data and d3 Java code [120] for generating the plot can be downloaded.

**Table 4.** Quantitative concordance between eBG and HierCC clustering. NOTE: *Salmonella*: Calculations were performed on 319,490 entries in EnteroBase which had been assigned to eBGs, HC900 and HC2000 clusters by December, 2021. The dataset only represented 411 eBGs, 690 HC900 clusters and 312 HC2000 clusters, and consists of a subset of all genomes because new eBGs have not been created in recent years. *Escherichia/Shigella*: HC1100 and HC2000 assignments were from 143,520 genomes which had been assigned to ST Complexes as well as to HC1100 and HC2350 clusters in December, 2021. At that time 186,852 genomes had been assigned to HC1100 and HC2000 clusters, with AMI of 0.59 and ARI of 0.2. *Streptococcus pneumoniae* consisted of GSPC assignments for 18,147 genomes from Supplemental Table 2 and CC assignments for 13,396 genomes from Supplemental Table 1 to Gladstone et al. [107] which were imported into User defined fields in EnteroBase as described [28]. ARI and AMI were calculated with the functions adjusted_rand_score() and adjusted_mutual_info_score() in the Python 3 library sklearn.metrics V1.0.1 [119].

|  |  |  |  |
| --- | --- | --- | --- |
| *Salmonella* |  |  |  |
| Test statistic | eBG-HC900 | eBG-HC2000 | HC900-HC2000 |
| Adjusted Mutual Information Score (AMI) | 0.985 | 0.948 | 0.942 |
| Adjusted Rand Index (ARI) | 0.982 | 0.856 | 0.866 |
| *Escherichia/Shigella* |  |  |  |
| Test statistic | ST Cplx-HC1100 | ST Cplx-HC2000 | HC1100-HC2000 |
| Adjusted Mutual Information Score (AMI) | 0.940 | 0.688 | 0.692 |
| Adjusted Rand Index (ARI) | 0.871 | 0.299 | 0.337 |
| *Streptococcus pneumoniae* |  |  |  |
| Test statistic | CC-HC100 | GSPC-HC100 | GSPC-HC160 |
| Adjusted Mutual Information Score (AMI) | 0.959 | 0.956 | 0.980 |
| Adjusted Rand Index (ARI) | 0.987 | 0.906 | 0.949 |

HC2000 Lineages with uniform O group included the Para C Lineage (HC2000_1272; serovars Paratyphi C, Typhisuis, Choleraesuis and Lomita [46]) which emerged about 3500 years ago (Supplemental Text) [46, 80]. The Para C Lineage is uniformly serogroup O:7, which is not particularly surprising because all its serovars have identical antigenic formulas, and are only distinguished by biotype. The Enteritidis Lineage (HC2000_12) consists of serovars Enteritidis (1,9,12:g,m:-), Gallinarum and its variant Pullorum (1,9,12:-:-), and Dublin (1,9,12:g,p:-:-) [45] (Fig. 5), and is uniform for O:9. The Typhimurium Lineage (HC2000_2) includes serovars Typhimurium, Heidelberg, Reading, Saintpaul, Haifa, Stanleyville and others [45]. This Lineage is also uniform for O group because all these serovars are O:4. The Mbandaka Lineage (HC2000_4) includes serovars Mbandaka (6,7,14: z10:e,n,z15) and Lubbock (6,7:g,m,s:e,n,z15), and is uniformly O:7. However, three Lineages were exceptional, and did contain more than one O serogroup. HC2000_299 is a mixture of O:4 (HC900_299; serovar Agama [28]) and O:7 (HC900_127074; serovar Nigeria). HC900 clusters within HC2000_54 are O:4 (HC900_54: Bredeney, Schwarzengrund; HC24937: Kimuenza) or O:9,46 (HC900_57: Give). Similarly, HC900 clusters within HC2000_106 are O:3,10; O:9; O:9,46; O:4: or O:8. These observations confirm and extend the prior conclusions about concordance between natural populations and serogroup within *Salmonella*.

#### Escherichia coli

Patterns of multilocus isoenzyme electrophoresis were used in the 1980s to provide an overview of the genetic diversity of *E. coli* (see overview by Chaudhuri and Henderson [89]). Those analyses yielded a representative collection of 72 isolates [90], the EcoR collection, whose deep phylogenetic branches were designated haplogroups A, B1, B2, C, D and E [91]. Several haplogroups have since been added [92, 93]. The presence or absence of several accessory genes can be used for the assignment of genomes to haplogroups with the Clermont scheme [94, 95], and the haplogroup can also be predicted *in silico* from genomic assemblies with ClermontTyping [96] or EZClermont [97], both of which are implemented within EnteroBase. However, the Clermont scheme ignores *Shigella*, which consists of *E. coli* clades despite its differing genus designation [26,42,43], and makes multiple discrepant assignments according to phylogenetic trees [28]. The Clermont scheme also does not properly handle the entire diversity of species and environmental clades/subspecies within the genus *Escherichia* [28,58,59,61].

EnteroBase HierCC automatically assigns genomes within the *Escherichia/Shigella* database to the cgMLST equivalents of ST Complexes (HC1100 clusters) and Lineages (HC2000 clusters) (Table 1). It also perpetuates the ST Complexes that were initially defined for legacy MLST by Wirth *et al.* [26]. However, the numbers and composition of legacy ST Complexes have not been updated since 2009 because additional sequencing data defined intermediate, recombinant genotypes that were equidistant to multiple ST Complexes, and threatened to merge existing ST Complexes. In contrast, HC1100 clusters do not merge: Intermediate genotypes are rare because cgMLST involves 2512 loci while legacy MLST was based on only seven. Secondly, new genotypes which are similar to and equidistant to multiple clusters do not trigger merging because HierCC arbitrarily assigns them to the oldest of the existing alternatives [32]. Finally, unlike ST Complex designations in legacy MLST, which have not been actively updated in *Escherichia* since 2006, HierCC clustering is an ongoing, automated process for all new genotypes.

Frequent recombination in *E. coli* results in poor bootstrap support for the deep branches in phylogenetic trees of concatenated genes [26], and we were unable to identify a unique HierCC level which was largely concordant with haplogroups according to the Clermont scheme. However, similar to *Salmonella,* HC1100 clusters identify clear population groups, which are highly concordant with legacy ST Complexes ( AMI = 0.94) (Table 4). Unlike *Salmonella*, HC2000 clusters did not in general mark recognizable additional population structure beyond that which was provided by ST Complexes, and HC2000 clusters are only moderately concordant with either legacy ST Complexes (AMI = 0.69) or their cgMLST equivalent, HC1100 clusters (AMI = 0.69) (Table 4). Multiple *Shigella* species are exceptions to this rule and their legacy ST Complexes [26] equate to cgMLST HC2000 clusters rather than HC1100 clusters according to phylogenetic trees. HC2000_305 replaces ST152 Cplx, and encompasses all *S. sonnei* (Fig. 6). HC2000_192 replaces ST245 Cplx, and contains many *S. flexneri*. However, *S. flexneri* O group F6 is in an HC1100 cluster together with *S. boydii* B2 and B4 within HC2000_1465 (ST243 Cplx). HC2000_1465 includes a second HC1100 cluster with *S. boydii* O groups B1 and B18, and a third with *S. dysenteriae* D3, D9 and D13 (Fig. 6). HC2000_4118 replaces the combination of ST250 Complex and ST149 Complex, which have merged, and contains both *S. dysenteriae* as well as *S. boydii*.

**Figure 6.**
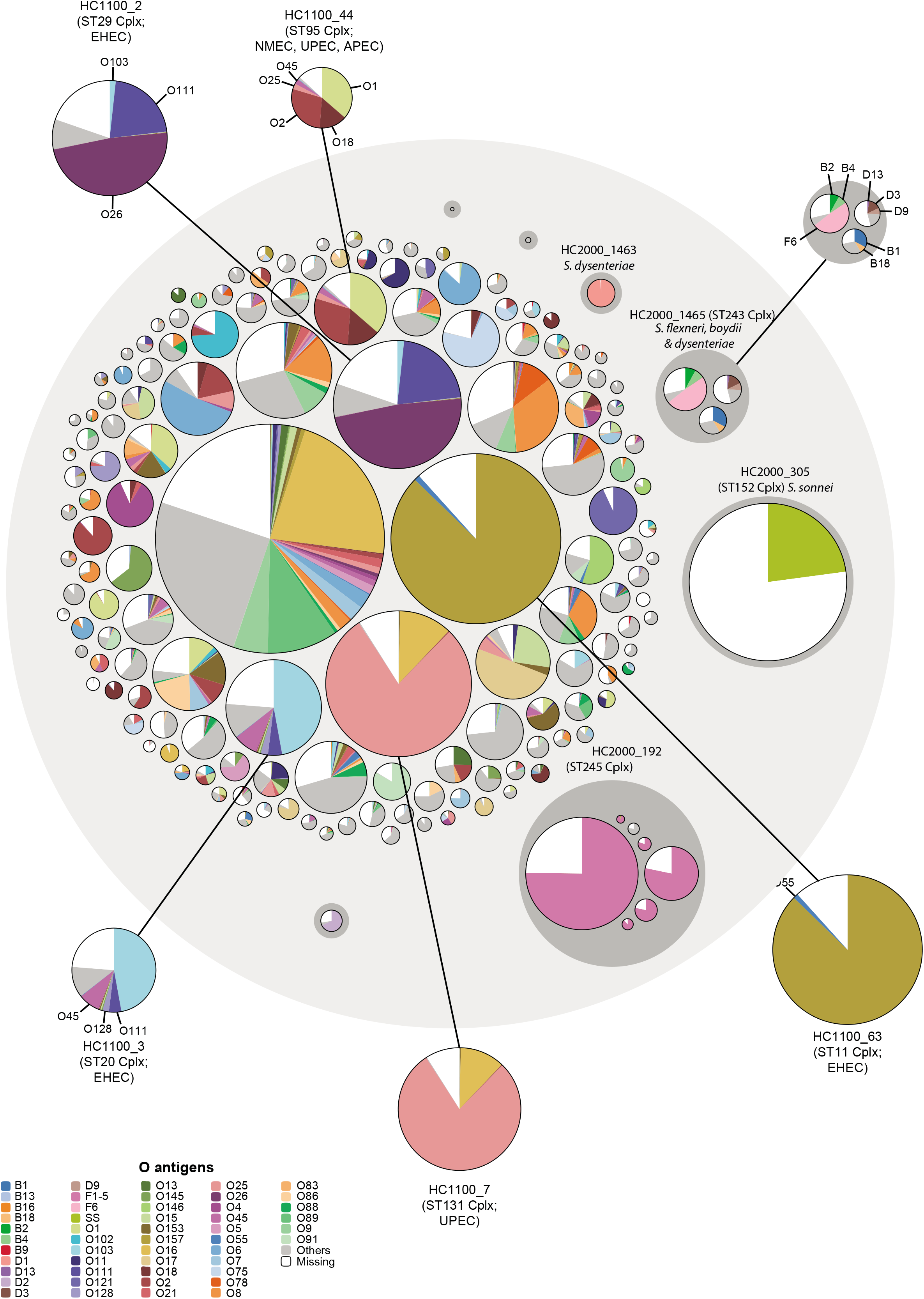
Hierarchical population structure of O serogroups in *Escherichia coli/Shigella*. Hierarchical bubble plot for 167,312 genomes of *Escherichia coli* or *Shigella* in HC2350_1 (large grey circle) that were available in EnteroBase in April, 2021. Seven HC2000 Lineages encompassing 15 HC1100/ST Complexes of *Shigella* are arbitrarily indicated at the right. HC2350_1 also includes 15 other *E. coli* HC2000 Lineages that encompass 144 other HC1100 clusters that each contains at least 50 *E. coli* genomes. The figure shows only the HC1100 clusters and not the corresponding HC1100 clusters. Numbers of genomes assigned to individual O serogroups (legend) are indicated by pie chart wedges within the HC1100 circles. Selected HC1100 clusters are drawn on larger scale with indications of phenotype and nomenclature outside the main circle, connected to the original circles by lines. An interactive version of this figure can be found at https://observablehq.com/@laurabaxter/escherichia-serovar-piechart, from which the representation, raw data and d3 Java code [103] for generating the plot can be downloaded.

Unlike *Salmonella*, O serogroups are remarkably variable within HC1100 clusters of *E. coli* (Fig. 6). HC1100_63 (ST11 Complex) combines classical EHEC strains of serovar O157:H7 as well as O55:H7 [98], and other, atypical ancestral EPEC isolates of serovar O55:H7 [99]. Multiple other HC1100 clusters that contain EHEC strains also encompass multiple O groups, including HC1100_2 (ST29 Cplx; O26, O103, O111) [100] and HC1100_3 (ST20 Cplx; O45, O103, O111, O128). Similarly, multiple HC1100 clusters of *E. coli* that cause extra-intestinal diseases also contain a variety of O groups, including HC1100_7 (ST131 Cplx; O16, O25) [23,101,102] and HC1100_44 (K1-encapsulated bacteria of the ST95 Cplx; O1, O2, O18, O25, O45) [26,103,104]. Indeed, the primary impression from Fig. 6 is that O group variation within an HC1100 cluster is nearly universal throughout *E. coli*, confirming prior conclusions about the high frequency of homologous recombination in this species [26, 92].

#### Other genera

Correlations between intra-specific HierCC clusters and population groupings at multiple levels, including ST Complexes and ribotypes have been described in *Clostridioides difficile* [105]. It remains to be tested whether HierCC can identify populations within *Yersinia* because bacterial populations in *Y. pseudotuberculosis* are largely obscured by recombination [106]. Similarly, we are not aware of extensive efforts to identify bacterial populations which could be compared with HierCC within *Vibrio*. However, recent work has described the assignment of large numbers of genomes of *Streptococcus pneumoniae* to legacy Clonal Complexes as well as GSPC (Global Pneumococcal Sequence Clusters) [107], initially with PopPunk [108] and more recently with Mandrake [109]. Most of those genomes have been assembled within EnteroBase, and assigned to HierCC clusters. We therefore compared the HierCC and GSPC assignments for 18,147 genomes, and also compared them to Clonal Complexes (CCs) based on legacy MLST (13,396 genomes). HC160 clusters yielded the highest AMI and ARI scores with GSPC clusters while CC was most concordant with HC100 clusters (Table 4). These conclusions were also supported by visual comparisons of the colour-coding of nodes within a Neighbor-Joining tree based on cgMLST distances according to the various clustering criteria (Supplemental Fig. S4). Thus, HC100 clusters seem to be concordant with Clonal Complexes within *S. pneumonia* and Mandrake/PopPunk clustering equates roughly with HC160 clusters.

## Discussion

In 2013, we anticipated that EnteroBase might eventually contain as many as 10,000 genomes of *Salmonella* and *Escherichia*. By the end of 2021 it hosted >600,000 assembled genomes of *Salmonella, Escherichia/Shigella, Streptococcus*, *Vibrio*, *Clostridioides* and *Yersinia*, and has become one of the primary global sources of their genomic data. An analyses of their genomic properties is facilitated by software tools (Table 2) that support indexing of genetic diversity, searching for genomes with specific metadata or genetic relations and investigating stable population structures at multiple levels. Here we focused on the automated identification and designation of species/subspecies and populations on the basis of HierCC of core genome MLST genotypes. The data demonstrate that HierCC can completely replace classical taxonomic methods for genomes of *Salmonella, Escherichia/Shigella, Vibrio*, *Clostridioides* and *Yersinia*, with only few minor exceptions. HierCC is also a complete replacement for legacy MLST regarding the assignments to populations within *Salmonella*, *Escherichia/Shigella* and *Streptococcus pneumoniae*.

### Taxonomic assignments

The paired ML super-trees based on core SNPs and presence/absence of accessory genes from each genus were generally concordant in their clustering patterns, indicating that those genera contain well defined and reproducible species/subspecies independent of the genetic criteria. Those phylogenetic clustering were then used to resolve the most approrpriate designations for genomes which were assigned contradictory designations by their taxonomic metadata, and clusters based on 95% ANI or HierCC. The frequency of agreement with current taxonomy was comparable for HierCC and 95% ANI at the species level within *Salmonella, Escherichia/Shigella, Vibrio*, *Clostridioides* and *Yersinia* but 95% ANI was generally unable to resolve sub-species whereas HierCC did not distinguish between sub-species and species clusters. Classical taxonomical designations exhibited multiple, glaring problems in each genus and we replaced those problematical designations in EnteroBase with labels based on the automated, HierCC-based species/subspecies taxonomy. We also defined novel species/sub-species, and labelled them with HierCC-based designations.

Why were there so many obvious discrepancies between the different approaches? One obvious reason is that we have discarded the medical tradition of retaining discrete species names for pathogens that cause particular diseases, and recommend designating *Y. pestis* as a sub-species of *Y. pseudotuberculosis.* Other discrepancies may reflect technical errors in manipulating data or uploading information to public databases, or simple strain mix-ups [41, 45]. Possibly the discrepancies were particularly obvious because the scale of our analyses were performed at an unprecedented scale over multiple genera. Many insights can be attributed to the facile ability to investigate large databases within a graphic framework that is provided by EnteroBase, and to our optimization of HierCC levels for each genus. However, we failed in a similar attempt to optimise ANI levels for recognizing both species and sub-species.

Our taxonomic changes in *Yersinia* are likely to be highly controversial. According to both ML trees and HierCC, the current designation “*Y. enterocolitica”* encompasses four, previously unnamed species/sub-species. These taxa are not distinguished by 95% ANI and had not previously been assigned distinct names. We maintain continuity with the traditional designation of *Y. enterocolitica*, and simply add HierCC affiliations to the species name, e.g. *Y. enterocolitica* HC1490_2. Alternatively, these clusters of strains could be considered to represent sub-species and their name modified slightly, e.g. *Y. enterocolitica* subspecies HC1490_2, similar to our downgrading of the named species *Y. pestis, Y. similis* [70] and *Y. wautersii* [69] on phylogenetic grounds to subspecies of *Y. pseudotuberculosis* (Fig. S1). Finally, we defined eight new species/sub-species within *Yersinia* that were not previously differentiated without submitting traditional evidence for simple phenotypic differences, and named them after their HC cluster, e.g. *Yersinia HC1490_4399*. We also followed comparable strategies for the other genera analyzed here.

In *Salmonella,* HierCC and 95% ANI identified a new species, *Salmonella HC2850_215890*, and a new subspecies, *S. enterica* subsp. HC2850_222931 (Fig. 1). Five new species were identified within *C. difficile* by HierCC as well as ANI, and HierCC identified two additional new sub-species. In *Escherichia* we provide HierCC designations for multiple clusters of bacteria from environmental sources near Hong Kong that were previously unnamed.

The ML trees presented here (Fig. S2) support prior definitions of high order clades containing multiple species in *Vibrio* on the basis of MLSA [72]. We also observed a general concordance between ANI and HierCC for 109 named *Vibrio* species, However, 43 HierCC clusters of species rank had not previously been properly classified with species designations, and 17 other species names were superfluous. These changes are fully described in Supplemental text, allowing closer scrutiny by the *Vibrio* taxonomic community for consistency with other properties.

HierCC was so effective in clarifying the taxonomies of these five species that we had hoped that it could also facilitate the taxonomic classification of *Streptococcus.* 102 HC363 clusters were indeed concordant with 95% ANI and taxonomic designations, with only minor exceptions. However, extensive discrepancies existed between 95% ANI and HierCC for numerous other clusters, and both approaches showed multiple discrepancies with classical taxonomic designations. The existence of multiple 95% ANI clusters within *S. mitis* and *S. oralis* and other species of *Streptococcus* has previously been commented on by Kilian and his colleagues [15,16,73,74]. Our observations extend these taxonomic problems to multiple additional species in which HC363 does not distinguish between pairs of named species (Fig. 4). Neither 95% ANI nor cgMLST HierCC seems to provide a suitable general strategy for elucidating the taxonomy of all of *Streptococcus*, and we failed to find a general solution to this problem.

### HierCC *versus* classical taxonomy

Classical microbiological species taxonomy involves identifying a type strain whose phenotype can be distinguished from all other type strains, identifying additional isolates with similar phenotypes, demonstrating distinct clustering from other known species in phylogenetic trees based on DNA or amino acid sequences, and publishing a report in one of a very limited number of acceptable journals. Such species definitions are then considered tentative until an international committee has approved them. Other forms of identifying species that include sole reliance on DNA sequence differences are not acceptable [7, 8]. The metagenomics community and scientists working with uncultivated organisms from the environment have largely liberated themselves from such regulations, and tend to use operational taxonomic units (OTUs) as taxonomic entities. However, the taxonomic species structure of many microbes from environmental sources remains fuzzy, or does not clearly correspond to classical taxonomy (e.g. *Prochlorococcus* [110] or *Synechococcus* [111]). Furthermore, we are not aware of any other method that able to assign 1000s of genomes per day to existing taxa and also reliably identify new taxa automatically as they appear. HierCC performs this task with bravura for the genera in Table 1, with the notable exception of *Streptococcus*.

We have taken the liberty of dropping all attempts to reconcile HierCC species/subspecies clusters with classical prescriptions for how to define a species. Instead we have adopted the practice of using HierCC cluster designations within EnteroBase for the nomenclatures for species/subspecies groupings which had not yet been identified by others, and eliminated from EnteroBase multiple species designations of type strains in *Vibrio* and *Streptococcus* which did not match the ML tree topologies and HierCC clusters. These actions provide a uniform base for the future additions of additional species and ensured that future genomes can be correctly assigned to uniform clusters of related strains. Scientists wishing to use these schemes to identify the species of their bacterial isolates can upload their sequenced genomes to EnteroBase. The HierCC assignments will be available within hours. We also welcome additional curators of these databases who are willing to test and improve the current taxonomic assignments we have implemented. But we reject the concept of designating our assignments as tentative until they are confirmed in several years by an international committee.

### HierCC and Populations

The first bacterial taxonomic designations were assigned over 100 years ago. Bacterial population genetics is much younger, and has many fewer practitioners. Initial population genetic analyses in the early 1980s subdivided multiple bacterial species into intra-specific lineages based on multilocus enzyme electrophoresis [90]. This methodology was replaced in the late 1990s by legacy MLST based on several housekeeping genes [21], which is currently being replaced by cgMLST based on all genes in the soft core genome [22, 30]. EnteroBase calculates genotypes for both types of MLST, as well as for rMLST [28]. Legacy STs differ from cgSTs due to different levels of resolution. However, the boundaries of ST Complexes/eBGs are highly concordant with HierCC clusters, with AMI indices of 0.985 for eBGs *versus* HC900 *in Salmonella* and 0.94 for ST Complexes *versus* HC1100 clusters in *Escherichia/Shigella* (Table 4). We recommended previously that serovars should be replaced by eBGs in *S. enterica* [27] and the data presented here show that HC900 clusters are an even better replacement. Our data also indicate that Lineages in *S. enterica* correspond to HC2000 clusters, and that these tend to be highly uniform for O group. The data presented here also indicate that HC1100 groups are a good replacement for detecting populations within *Escherichia* as are HC100 clusters for CCs in *S. pneumoniae* [107] (Table 4).

We interpret these consistencies between legacy MLST and cgMLST as reflecting the existence of natural populations. We previously claimed that legacy ST Complexes in *E. coli* were unstable due to frequent recombination [26], and were therefore not surprised at their tendency to merge as additional isolates were sequenced. However, legacy MLST is based on only seven genes. The high resolution of cgMLST and the decision to assign new genotypes that are equidistant from multiple HierCC clusters to the oldest cluster have largely negated these problems, and *E. coli* HC1100 clusters correspond to bacterial groupings that have been independently identified by multiple phenotypic patterns. In the early 1980s, MA began his academic research on bacterial pathogens with *E. coli* that expressed the K1 polysaccharide capsule [103]. He observed unusually uniform patterns of electrophoretic mobility of major outer membrane proteins across multiple isolates, and interpreted those bacteria as representing “clones”. These “clones” correspond to HC1100 clusters, including HC1100_44 (ST95 Complex) which continues to cause invasive disease in humans and animals around the globe [104]. The 1980s analyses showed that these K1 bacteria were variably O1, O2, or O18, and that the serotype variants differed both in their invasiveness and in the hosts which they infected [112]. As indicated in Fig. 6, HC1100_44 includes O groups O25 and O45 in addition to O1, O2 and O18. Similarly, HC1100_7 (ST131) represents another major cause of extra-intestinal disease in humans [23,101,102]. EHEC bacteria are notorious for their association with hemolytic uremic syndrome (HUS). Many of them belong to distinctive HC1100 clusters, including O157:H7/O55:H7 (HC1100_63, ST11 Cplx [99, 113]) and O26, O103 and O111 EHEC bacteria (HC1100_2, ST29 Cplx [100]) (Fig. 5).

In contrast to these correlations with epidemiological groupings, we were unable to identify any other HierCC levels that were consistently concordant with other intra-specific phylogenetic subdivisions based on SNP trees, including haplogroups [91] or Clermont typing [94, 95] in *E. coli* [28] and Clades [76–79], lineages [75] or Branches [80] in *S. enterica.* Instead, we found that a subdivision we refer to as Lineages are marked by some HC2000 clusters in *Salmonella* and *Escherichia/Shigella*.

### Lineages

In *E. coli*, HC2000 Lineages were particularly appropriate for defining the clustering level of seven groups of *Shigella* genomes (Fig. 6). Lineage designations did not add additional insights into other populations because were almost entirely encompassed by HC1100 clusters or occasionally even lower level clusters, such as HC400. In *S. enterica* subspecies *enterica*, multiple HC2000 Lineages corresponded to genetically related combinations of multiple HC900 clusters, in some cases with distinctive serovar designations. However, all but three of the major Lineages were predominantly uniform in O group. The three exceptional HC2000 Lineages (HC2000_54, HC2000_106, HC2000_299) might be worth exploring in greater detail to reconstruct the recombination that has resulted in their differing LPS epitopes.

HC160 clusters, the Lineage equivalent in *S. pneumonia,* were strongly concordant with PopPunk GSPC clustering. Otherwise, little is yet known about the general properties of Lineages in other genera, except that their properties are likely to vary with species or genus. For example, *S. enterica* HC2000 Lineages were uniform for O group whereas O serogroups were already heterogeneous within HC1100 clusters/ST Complexes within *E. coli* (Fig. 6).

### Future prospects

EnteroBase was conceived to satisfy a need that MA and ZZ perceived in 2014 [1]. We contend that it is now fit for purpose for investigations of multiple bacterial genera by scientists ranging from beginners through to experts in the areas of microbial epidemiology and population genetics. Its original creators have now all left this project, but EnteroBase is being maintained as a service for the global community by the University of Warwick. Maintenance of its technical functions and databases are thereby assured for the near future. Further functional developments will, however, depend on increased participation and perception of ownership by its users. We perceive a general trend to focus on insular solutions that can satisfy the demands of individual bioinformaticians and regional diagnostic laboratories. Such approaches can yield relatively rapid progress in solving short-term needs. However, a global overview of genomic diversity needs central databases to ensure definitive terminology. Pooling efforts on a central endeavor at the scale represented by EnteroBase would ensure that it continues to function over decades, is representative over all continents, and serves the global community even better. We therefore welcome additional curators and scientific experts as well as bioinformatics collaborations to help improve EnteroBase even more.

## Methods (see Supplemental Text)

### Funding

MA was supported by grant 202792/Z/16/Z from the Wellcome Trust (https://wellcome.org/).

The funders had no role in study design, data collection and analysis, decision to publish, or preparation of the manuscript.

### Code

cgMLSA is available for downloading at https://github.com/zheminzhou/cgMLSA. All trees in figures and genomic data are freely available within EnteroBase for interactive examination. Other code is described in Supplemental Text.

### Data

Datasets used for the analyses presented here have been deposited for permanent public access at http://wrap.warwick.ac.uk/162247/

## Acknowledgements

We are very grateful to François-Xavier Weill for permission to include Table S8 for public access in Supplemental Material. We are very grateful to François-Xavier Weill and Tandy Warnow for their comments on an earlier draft of this manuscript.

