## Supplemental Text for "EnteroBase: Hierarchical clustering of 100,000s of bacterial genomes into species/sub-species and populations"

Supplementary Text

### Correspondence between species designations and ANI or HierCC

Multiple short read sequences uploaded to EnteroBase from public depositories such as ENA or by EnteroBase users lack information on species, or contain incorrect species assignments. On irregular occasions, the EnteroBase databases have been intensively curated to provide/correct species assignments within the metadata. Prior work focused on *Salmonella* and *Escherichia.* As part of the preparations for this publication, species metadata were curated in late 2021 for the genera *Clostridioides, Streptococcus, Yersinia* and *Vibrio.* The existing metadata was subjected to detailed comparisons between prior species designations and the branching patterns within Maximum Likelihood phylogenetic super-trees of representative collections of genomes from EnteroBase (Table 3) that were calculated from the presence/absence of accessory genes or from the nucleotide variants in non-repetitive regions within core genes. Where available, genomes from all type strains for individual species were also included in the genomic collections, and that status listed in the Comments field. The accessory genome was calculated as the variable genes within a pan-genome generated for that representative collection with PEPPAN [1], and presence/absence data was calculated from the individual genomes with EToKi [2]. SNPs in core genes were also calculated with EToKi. Distance matrices and super-trees were calculated with cgMLSA (see Methods below). All resulting pan-genomes and the distance matrices have been deposited at the University of Warwick WRAP collection for permanent storage, which can be accessed at http://wrap.warwick.ac.uk/162247/. The main manuscript presents some of this data for *Salmonella*, *Escherichia/Shigella*, *Clostridioides* and *Streptococcus*. Here we focus on *Yersinia* and *Vibrio.* URLs to Interactive trees are listed in the Figure legends and https://enterobase.readthedocs.io/en/latest/species_trees.html.

#### *Yersinia*

*Yersinia* contains multiple pathogenic and non-pathogenic species. Reuter *et al.* [3] published phylogenetic analyses in 2014 of SNPs in concatenated sequences of 84 housekeeping genes from 241 genomes representing the known taxonomic diversity of *Yersinia* at that time. A subsequent analysis in 2019 by Savin *et al.* [4] summarised data for a core genome MLST scheme of 500 genes from 1,346 genomes. Savin *et al*. noted that genomic data in the public domain were often associated with faulty taxonomic designations, and they also identified seven additional species-like clades, which were subsequently given species status [5,6]. These two publications indicate that the taxonomic designations of *Yersinia* are incoherent across the genus, and that no automated system exists for the recognition of novel species-level taxa as additional genomes are sequenced, or for the automated assignment of taxon-level designations. We therefore erxamined ML super-trees from the presence/absence of accessory genes, and of core non-repetitive SNPs from 1847 genomes that are representative of all *Yersinia* in EnteroBase. The topologies of these trees were comparable independent of the distance metric (Fig. S1 top *vs* bottom), and the phylogenetic clustering was also coherent with clustering according to 95% ANI (Fig. S1 A, C) *versus* HC1490 (Fig. S1B, D). However, they were partially incoherent with current taxonomic designations.

Our conclusions are summarised in Fig. S1. The clades containing genomes from type species were designated according to the corresponding taxonomic designations. Once the associations between HierCC clustering and taxonomic designations had been clarified, we changed the metadata in EnteroBase for each HC1490 cluster to correspond to the designations in Fig. S1. Where no type strain was known for an HC cluster, we used the cluster number as an equivalent to a species designation, and thereby defined eight novel species/sub-species designations within *Yersinia* according to their unique HierCC clusters: *Y. HC1490_419, Y. HC1490_435, Y. HC1490_440, Y. HC1490_449, Y. HC1490_457, Y. HC1490_1007, Y. HC1490_4399* and *Y. HC1490_4429*. For example, *Y. HC1490_435* and *Y. HC1490_449* for sister clades of *Y. mollareti.* These implementations provide an automated system for detecting and naming of new species/subspecies. However, similar to the results for *Salmonella* and *Clostridoides*, they also highlight some discrepancies between ANI 95% clusters and HC1490 clusters, which are indicated by asterisks in Figs. S1A and S1C. For example, ANI95% does not distinguish between *Y. HC1490_435, Y. HC1490_449* and *Y. mollaretti.* Similarly, ANI95% does not distinguish between *Y. pekkanenii* and *Y. HC1490_1007*, or *Y. alsatica* and *Y. HC1490_4429* or between *Y. occitanica* and *Y. thracica*. However, all other taxonomic designations were consistent according to both 95% ANI and HC1490 clustering except for *Y. enterocolitica* and the *Y. pseudotuberculosis* Complex.

According to Reuter *et al.* [3], *Y. enterocolitica* consists of four phylogenetic clades with different metabolic biotypes. Two 1A biotype clades within *Y. enterocolitica* are basal, and might have been ancestral. They lack the pYV virulence plasmid, which was likely acquired independently on two occasions during the subsequent differentiation of the 1B biotype clade and the 2-5 biotypes clade. These four groups also formed distinct phylogenetic branches in trees by Savin *et al.* [4]. Although they are indistinguishable by 95% ANI (Fig. S1A, C), all four clades correspond to distinct clusters according to HierCC HC1490 (Fig. S1B, D). These four clades therefore warrant being assigned to distinct species/sub-species, and have been renamed in EnteroBase. *Y. enterocolitica* HC1490_73 and *Y. enterocolitica* HC1490_764 designate the two pYV-less and avirulent 1A biotype clades. The highly virulent 1B biotype clade was designated *Y. enterocolitica* HC1490_2 and the moderately virulent clade of biotypes 2-5 is *Y. enterocolitica* HC1490_10. These names maintain the traditional species designation of *Y. enterocolitica* in order to promote a ready association with historical designations but also include a species specific HC1490 cluster designation.

In 1980, Bercovier *et al.* recommended designating *Y. pestis* as a sub-species of *Y. pseudotuberculosis* because their DND-DNA hybridisation values were so similar [7]. In 1999, Achtman *et al.* [8] showed that *Y. pestis* was a clade of the *Y. pseudotuberculosis* Complex [9] whose taxonomic name was retained because only *Y. pestis* causes bubonic plague. The *Y. pseudotuberculosis* Complex encompasses the gastrointestinal pathogen *Y. pseudotuberculosis*, and two additional distinct clades that have been given species status: *Y. similis* by Sprague *et al.* [10] and *Y. wautersii* (formerly the Korean group) by Savin *et al*. [11]. However, all these clades are in the same ANI95% cluster (Fig. S1A, C) and the same HC1490 cluster (Fig. S1B, D). Both the presence/absence tree (Fig. S1, top) and the SNP tree (Fig. S1, bottom) confirm that *Y. pestis* is nothing more than a phylogenetic clade of *Y. pseudotuberculosis*. Thus the *Y. pseudotuberculosis* Complex is not subdivided into the supposed four species by any of the neutral phylogenetic approaches. The different diseases induced by *Y. pestis* likely reflects the historical acquisition of distinctive plasmids [12] a few thousand years ago [13]. Species assignments in EnteroBase are based on HierCC, which ignores genes on variable plasmids. Therefore, in order to provide a uniform taxonomy of all *Yersinia*, it was necessary to downgrade the taxonomy of the supposed species *Y. pestis, Y. similis* and *Y. wautersii* to subspecies called *Y. pseudotuberculosis* subsp. *pestis, Y. pseudotuberculosis* subsp. *similis*, and *Y. pseudotuberculosis* subsp. *wautersii*, respectively. In order to avoid additional confusion, we propose referring to *Y. pseudotuberculosis* as *Y. pseudotuberculosis sensu strictu.*

***Vibrio***

We identified 152 HC1090 clusters within *Vibrio,* of which 133 were equivalent to 95% ANI clusters (Supplemental Table S1). Eleven HC1090 clusters each corresponded to between two and five 95% ANI clusters (Supplemental Table S2) and three 95% ANI clusters each corresponded to two to three HC1090 clusters (Supplemental Table S3). 119 HC1090 clusters contained at least one genome from a type strain but some contained type strains from more than one taxonomic unit. The species name assigned to the ANI cluster was that of the type strain for 101 HC1090 clusters with only one type strain and of the dominant species name for 18 HC1090 clusters without a type strain but in which the species name was consistent across multiple genomes. For 123 genomes in 43 other clusters lacking a type strain, or in which there were unresolvable contradictions, we assigned the HC1090 cluster number as a species designator. Examples of contradictions and solutions: 1) HC1090_1705 contained 2 genomes which had been isolated from marine surface water near Florida in 2016 whose species was specified as *V. parahaemolyticus.* However, *V. parahaemolyticus* corresponds to HC1090_5, and is not closely related to HC1090_1705. The two genomes were assigned the species designation *Vibrio* HC1090_1705, and *V. parahaemolyticus* was stored as the prior designation in the Comments metadata field. 2) the species designation *V. hepatarius* was assigned to genomes in HC1090_5006 because it includes its type strain, DSM 19134. A different genome in HC1090_2595 was also designated as *V. hepatarius* but its 95% ANI cluster and HC1090 cluster are distinct from the first genome, although on the same phylogenetic branch. The species name of the second genome was changed to *Vibrio* HC1090_2595 (Table S4). Similarly, the species name was retained for *V. gazogenes* type strain DSM 21264 (NCBI Biosample SAMN02745781) in HC1090_4486 whereas the species of a second supposed *V. gazogenes* strain, ATCC 43942 (NCBI Biosample SAMN06053751) in the distinct but related HC1090 and ANI clusters was changed to *Vibrio* HC1090_7553 (Table S4).

These curation steps resulted in species designations or their equivalent for 152 HC1090 clusters, and those names were applied to 9133 *Vibrio* genomes in those HC1090 clusters. These curation efforts corrected numerous mistakes in species designations within the public domain, including those resulting from the tendency of NCBI to use a strain name for a species name when no species name is specified. Unfortunately, some ANI clusters also included multiple species names that could not be resolved by this approach, and these 'dead species' (Tables S1 and S2) are currently not used in EnteroBase except in the

Comments field. They consist of the *Vibrio* species *albensis, antiquarius, atlanticus, barjaei, celticus, chemaguriensis, communis, coralliirubri, diabolicus, hyugaensis, inhibens, kanaloae, ordalii, qinghaiensis, shilonii, tasmaniensis, toranzoniae* (Table S4). In some cases it might be possible to reinstate one or more of the species names which were eliminated. One example is HC1090_1339 which initially contained both *V. atlanticus* and *V. tasmaniensis* designations in NCBI. These designations were assigned to genomes in a manner that was so inconsistent with phylogenetic trees that it was impossible to decide between possible alternatives purely on phylogenetic grounds. In contrast, HC1090_2627 is phylogenetically consistent internally with *V. kanaloae* being restricted to one sub-cluster and *V. toranzoniae* to another, but both are included within a single HC1090/ANI cluster. In this case, a taxonomic decision between one of the names would be an acceptable alternative. Similarly, HC1090_180 consists of two 95% ANI clusters, one largely consisting of *V. jasicida* and the other of *V. hyugaensis*

**Historical reconstructions.**

In September 2021, the 12 HC2850 clusters corresponding to species/subspecies of *Salmonella* encompassed 2185 HC2000 Lineages (Table 1). However, their age, phylogenetic relationships and natural history largely remain to be elucidated except for the Para C Lineage (HC2000_1272) [14] and the AESB branch [15], which have recently been demonstrated to have caused human infections over millennia. The Para C Lineage includes serovar Paratyphi C (HC900_4381) which only infects humans. This Lineage also encompasses serovar Choleraesuis (HC900_1272 & HC900_5875), a swine pathogen that can infect humans, and Typhisuis (HC900_12567 & HC900_18096), which causes a typhoid-like disease in swine [14]. Although they are designated as distinct serovars, all three have the same antigenic formula, 6,7:c:1,5, and are differentiated by metabolic properties. Genomes reconstructed from ancient human tooth pulp identified Paratyphi C from historical human infections in Europe between 1200 and 1652 CE (Norway [14], Germany [16] and Spain [17]) and Mexico in 1545 CE [18]. Serovar Choleraesuis caused human infections in Europe about 1700 years ago, and the Para C Lineage split into multiple HC900 clusters from its common ancestor (tMRCA) 3500 years ago [14,15,17,18]. Recently, two ancient genomes within the Para C Lineage were reconstructed from DNA preserved in human tooth pulp from 3000 year old skeletons in XinJiang, China [19]. These mapped to an early, now extinct branch. The Para C Lineage also includes the rare serovar Lomita (human infections; HC900_10760), which defines an even older side branch [14]. Including Lomita, the tMRCA of the entire Para C Lineage was estimated as 4300 – 12,000 years by Key *et al.* [15]. Key *et al.* also reconstructed still other *Salmonella* genomes from human dental pulp from multiple other Neolithic infections. These were assigned to still deeper branches within an “Ancient Eurasian Super Branch (AESB)” [15], whose functional and phylogenetic relationships to modern genomes have not yet been fully elucidated.

**Sample bias.**

Genomes in the public domain have an enormous sample bias towards bacteria from human disease and food-borne infections in the U.S.A. and the U.K. For example, of 312,196 *Salmonella* genomes in EnteroBase, 44% (137,455) originated in the U.S.A. and 22% (68,104) in the U.K. 92% of the U.S.A. genomes were uploaded by the Centers for Disease Control, PulseNet, Food & Drug Administration or US Department of Agriculture, and 89% of the U.K. genomes had been uploaded by Public Health England. However, EnteroBase offers the same automated pipelines to its 4,800 users who also upload short reads plus metadata. As a result, 14% (44,041) of the *Salmonella* entries in EnteroBase had been uploaded by users. The vast majority of those sequences were from human salmonellosis in France (>26,000 uploads by the Institut Pasteur, Paris) or Scotland (>6,000 uploads by Scottish Microbiology Reference Laboratories, Glasgow). However, large numbers of sequence reads are being uploaded to EnteroBase from around the globe, and the initial sample bias is becoming less extreme. For example, 10,000 *Salmonella* genomes were from the UoWUCC project [20] which sequenced historical isolates from multiple sources, hosts and countries, and 4,208 other genomes were from the 10KSG project to sequence *Salmonella* from extra-intestinal diseases of humans in Africa and South America [21].

**Methods**

**Collections of representative genome sequences.** *Escherichia* (EcoRPlus) was described previously [2]. Other collections of representative genomes from six genera (Table 3) were selected in Jan 2021 using the following criteria. 1) All genomes derived from type strains for taxonomy where possible. 2) a single genome with the fewest uncalled nucleotides (Ns) from all other HC5 clusters (*Vibrio*, *Yersinia* and *Clostridioides*) or all other HC5 clusters with at least 5 genomes (*Salmonella* and *Streptococcus*). 3) one genome for each unrepresented HC400 (*Salmonella*) or HC50 (*Streptococcus*) cluster. *Escherichia* reps consisted of one representative genome from each of the 160 most common HC1100 clusters in HC2350_1 plus all genomes from other HC2350 clusters that were available in EnteroBase as of November 2020.

**ML super-trees.** For each representative genome collection, allelic sequences for that genus were retrieved from EnteroBase for each core gene in the cgMLST scheme. The sequences for each core gene were aligned with MAFFT [22], run with the option "—auto", and a gene tree of each of those alignments was calculated with RAxML-NG [23] , with the options '—tree pars{3} --model GTR+G'. The following steps were performed by the cgMLSA pipeline for super-tree construction, which can be downloaded from https://github.com/zheminzhou/cgMLSA. That pipeline performs the following sequential steps, as summarised in Supplemental Fig. S5.

**ASTRID guide tree.** A guide tree summarising the entire set of gene trees for each representative genome collection was created with ASTRID [24], and then evaluated with ASTRAL [25]. In brief, local quartet frequencies were calculated for each branch in the guide tree by comparing the topology of that guide tree with the topologies of all gene trees. Branches in the guide tree whose topologies were supported by at least 20% more local quartets than any alternative topology were scored as highly supported. The guide tree was split at each highly supported branch, and an artificial tip was appended to each resulting subtree to provide a permanent identifier (marker) for later re-joining to the subtree that had been removed. The result of these operations was a set of small and disjoint subtrees $S_i$, each of which contains a subset of genomes whose topologies were poorly supported in the guide tree plus one or more artificial tip markers that linked them to a other subsets *Sj via* a highly supported branch (Fig. S5C).

We then split each of the core gene trees into the corresponding subsets of genomes for each subtree using the Python ETE library [26]. All genomes within the subset corresponding to *Si* were labelled with a unique identifier (Fig. S5D) as was a random-genome from the core genome subset corresponding to *Sj.* The optimal topology of each subset was calculated again with RAxML-NG and ASTRAL was used to summarize the extracted gene subtrees from all core genes into sub-super-trees (Fig. S5E). A final complete super-tree was then constructed by connecting the sub-super-trees at the positions of the artificial tips from Fig. S5C. The local posterior branch support of the super-tree was calculated for all gene trees with ASTRAL, and Branch lengths in the super-tree were estimated using ERaBLE [27].

**Frequencies of O groups in *E. coli* and *Salmonella*.**

The hierarchical organization of HierCC data at the species/subspecies, Lineage and ST Complex/eBG levels was visualised with circle packing plots, showing HC2850, HC2000 and HC900 clusters for *Salmonella* and HC2350, HC2000 and HC1100 clusters for *Escherichia*. Circle packing (also called circular treemap) is equivalent to a treemap or dendrogram, where each node of the tree is represented as a circle and its sub-nodes are represented as internal nested circles (https://observablehq.com/@d3/circle-packing). Plots were created within ObservableHQ (https://observablehq.com/), a platform for data visualisation built on the D3 JavaScript library [28], using their integrated discovery environments or "notebooks".

The HierCC and serotype data for plotting were downloaded from EnteroBase in May 2020 for 167,297 *E. coli* strains and 274,824 *Salmonella* strains. We first extracted the serovars from all 790 *Salmonella* HC900 clusters which contained at least five entries in EnteroBase from the metadata Serovar field and the serovar predictions by SISTR 1.1.1 and SeqSero2. We then extracted the predominant consensus O serovar typical of ≥80% of entries for each source. For many large HC900 clusters, the three data sources were congruent. Where one of the three sources contradicted the two others, we accepted the majority serovar that was common to two data sources. In other cases without a consensus, the entries were manually examined within EnteroBase and with the help of GrapeTree plots of allelic distances. Patently wrong serovar metadata were deleted unless they had been deposited by the WHO Reference Laboratory at the Institut Pasteur, Paris and corresponded to genomes from a reference strain for individual, rare serovars. In almost all cases, manual inspection resolved the contradictions, empty metadata fields were updated to reflect the most reliable of the software predictions, and O groups were extracted from the consensus serovars according to Supplemental Table S8. Rare exceptional HC900 clusters which could not be reliably assigned serovar designations were not further modified. Fig. 5 summarizes the assignments of the most common 47 O serogroups to HierCC. The code and data are available at https://observablehq.com/@laurabaxter/salmonella-hierarchical-clusters.

For *E. coli*, O antigens were according to the predictions by *EBEis* within EnteroBase [2], and Fig. 6 shows the predicted frequencies of the 50 most common O antigens as pie charts with unique colour codes for each O antigen. The code and data are available at https://observablehq.com/@laurabaxter/escherichia-serovar-piechart.

17. de-Dios, T., Carrion, P., Olalde, I., Llovera, N. L., Lizano, E., Pamies, D., Marques-Bonet, T., Balloux, F., van Dorp, L., and Lalueza-Fox, C. 2021 *Salmonella enterica* from a soldier from

the 1652 siege of Barcelona (Spain) supports historical transatlantic epidemic contacts. *iScience* **24**, 103021. (doi:10.1016/j.isci.2021.103021 [doi];S2589-0042(21)00989-5 [pii])
