## Supplementary material for "EnteroBase: Hierarchical clustering of 100,000s of bacterial genomes into species/sub-species and populations": Figure S1

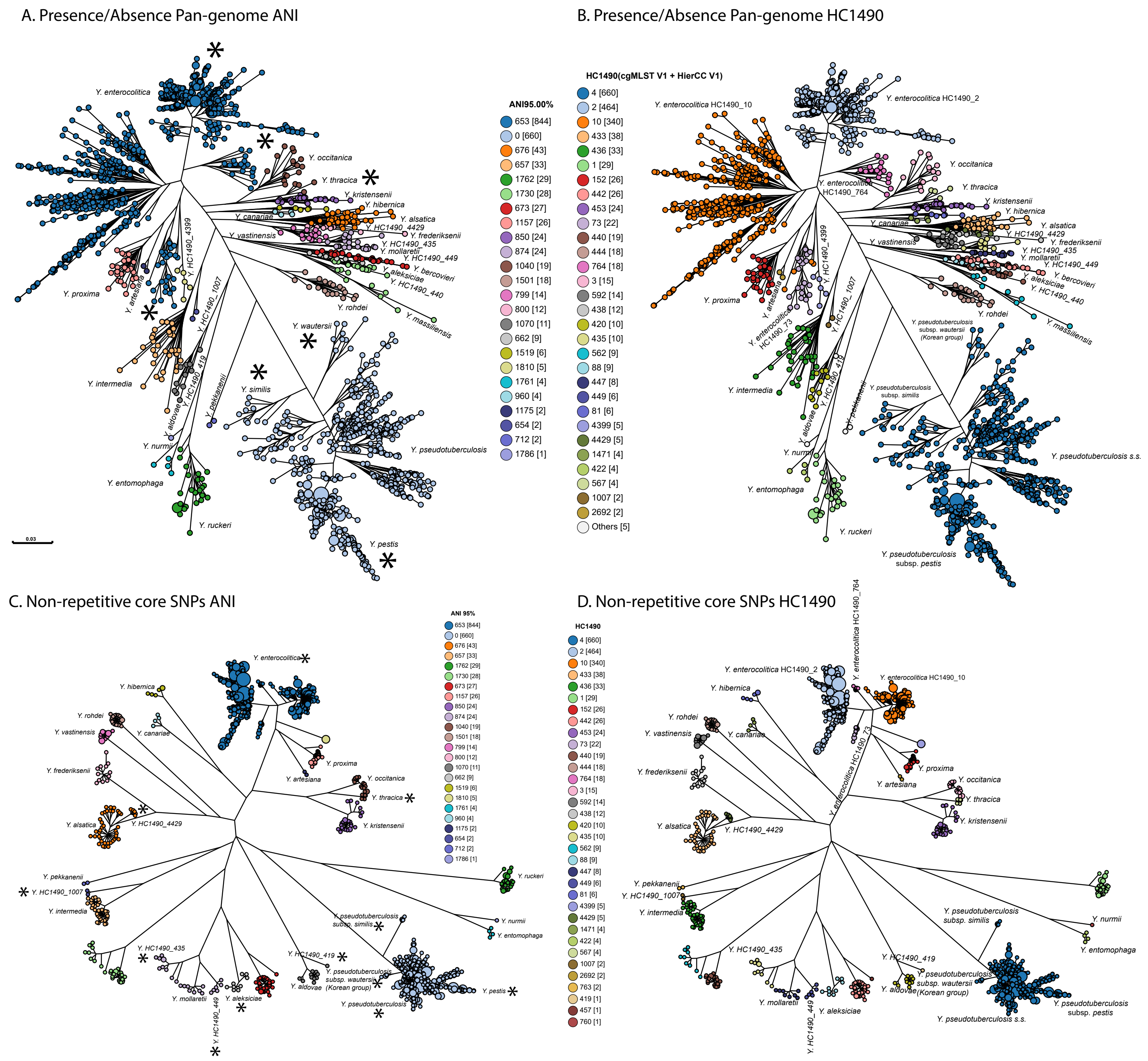


**Supplemental Figure S1.** Species and sub-species assignments within *Yersinia* with HierCC and ANI. A,B). Presence/Absence ML tree of 18,799 accessory genes among 1847 representative genomes of *Yersinia*, color-coded by 95% ANI cluster (A) or HC1490 cluster (B). C,D). SNP ML super-tree of 656,143 SNPs among 1553 soft core genes from 1847 representative genomes of *Yersinia*. Color-coded by 95% ANI cluster (C) or HC1490 cluster (D). The most common color codes are indicated in legends next to each plot. Within each tree, species designations are indicated next to the appropriate clusters. Note that the designations of *Y. enterocolitica* reflect its split into four species by HC1490 cluster which is not supported by 95% ANI clustering. Similarly, *'s in A and C indicate sub-species *similis*, *wautersii* and *pestis* of *Y. pseudotuberculosis* that were previously assigned distinct species designations. The novel *Yersinia* species *HC1490_1007*, *HC1490_440*, *HC1490_435*, and *HC1490_449* are also included in the figure. Further details can be found in Supplemental Text and interactive version of the GrapeTree renditions can be found at https://enterobase.warwick.ac.uk/ms_tree?tree_id=53270 (A,B) and https://enterobase.warwick.ac.uk/ms_tree?tree_id=53269 (C,D).
