## Supplementary material for "EnteroBase: Hierarchical clustering of 100,000s of bacterial genomes into species/sub-species and populations": Figure S2

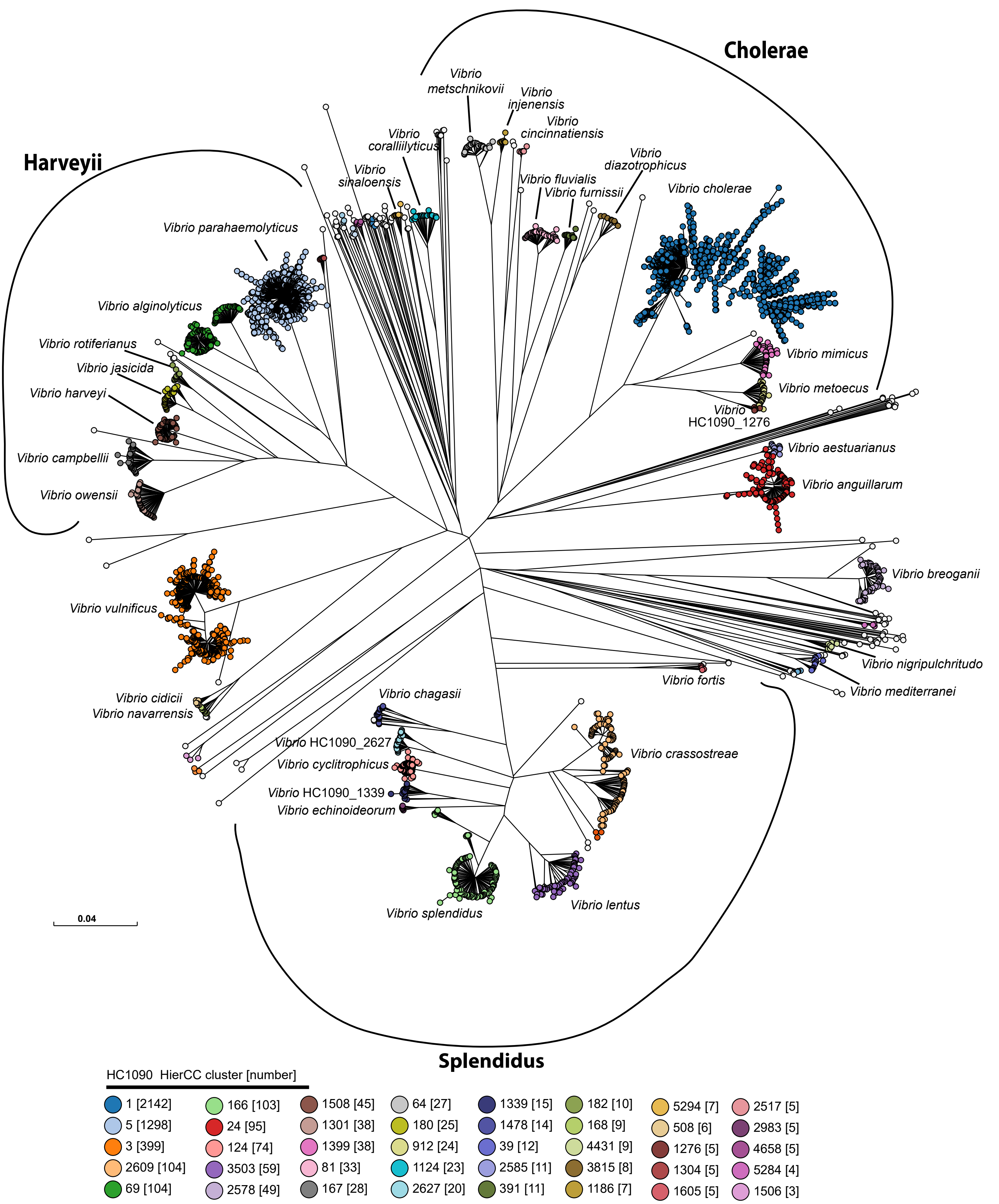


**Supplemental Figure S2.** Species and sub-species assignments based on HierCC within *Vibrio*. ML SNP super-tree of 776, 439 SNPs among 1128 soft core genes from 5032 representative genomes of *Vibrio*. Color-coded by HC1090 cluster. The most common color codes are indicated in the legend at the bottom. Species designations for the largest HC1090 clusters are indicated next to those clusters. Three higher order groups of species, Cholerae, Harveyii and Splendidus, that were defined by MLSA by Gomez-Gil et al., 2014 are indicated by arcs around the figure. Further details can be found in Supplemental Text and interactive versions of the GrapeTree renditions can be found at https://enterobase.warwick.ac.uk/ms_tree?tree_id=53265 (SNP tree) and https://enterobase.warwick.ac.uk/ms_tree?tree_id=53266 (presence/absence tree).
