## Supplementary material for "EnteroBase: Hierarchical clustering of 100,000s of bacterial genomes into species/sub-species and populations": Figure S3

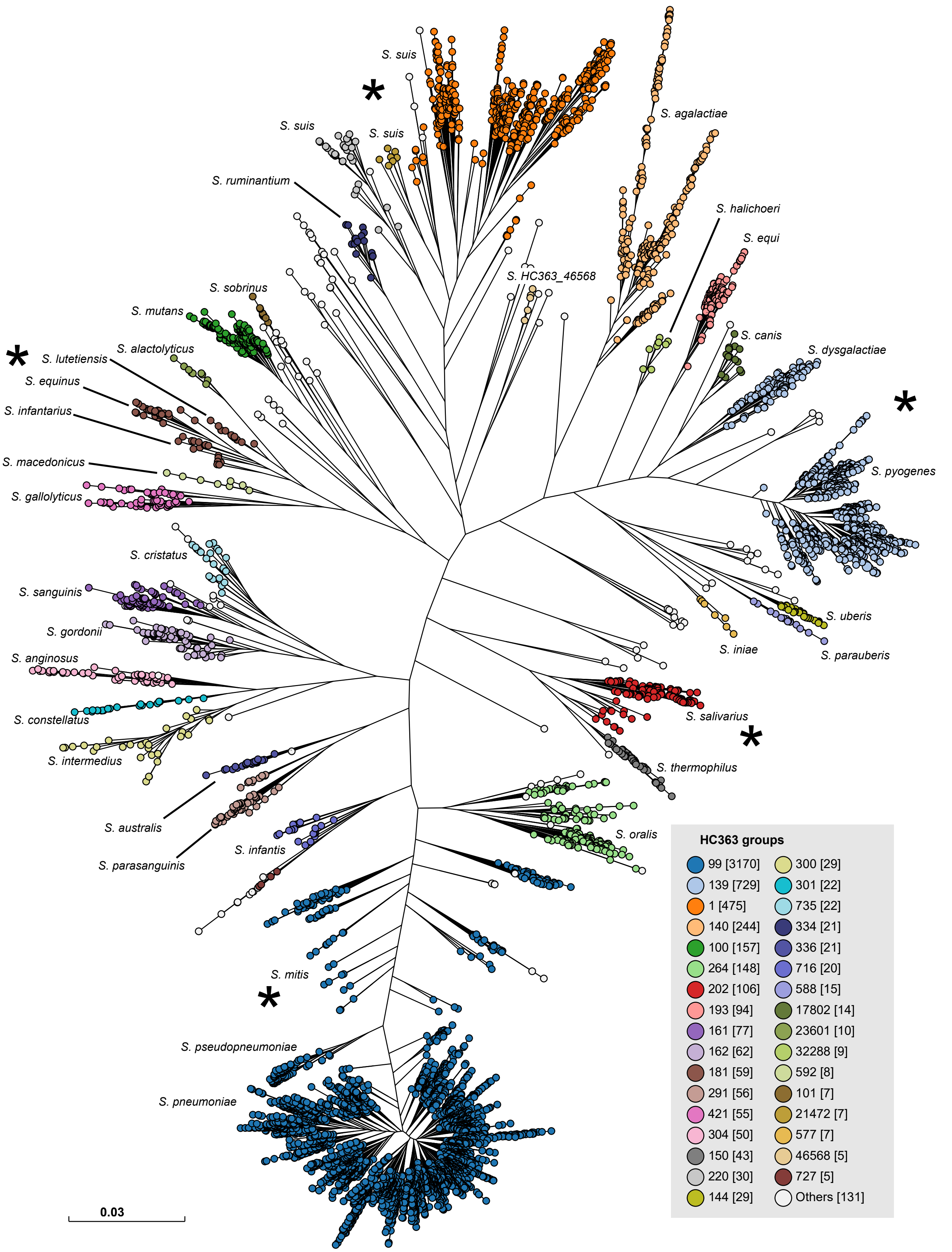


**Supplemental Figure S3.** Comparison of HC363 clusters with taxonomic designations in *Streptococcus* in an ML super-tree of the presence/absence of 31,630 accessory genes from 5937 representative *Streptococcus* genomes. Species names are indicated next to the phylogenetic clusters for genomes which mapped together with type strains, and according to published metadata. Nodes are colored by HC363 clusters, and exceptional assignments are indicated by asterisks next to *S. mitis*, which could not be distinguished by HC363 clustering from *S. pneumoniae* or *S. pseudopneumo-niae*, multiple phylogenetic and HierCC clusters within *S. suis*; *S. salivarius* and *S. vestibularis,* which were both HC353_202; *S. lutetiensis* and *S. equinus*, which were both HC363_181; and *S. dysgalactiae* and *S. pyogenes*, which were both HC363_139. An interactive version of this GrapeTree rendition can be found at https://enterobase.warwick.ac.uk/ms_tree?tree_id=53262.
