## Supplementary material for "EnteroBase: Hierarchical clustering of 100,000s of bacterial genomes into species/sub-species and populations": Figure S4

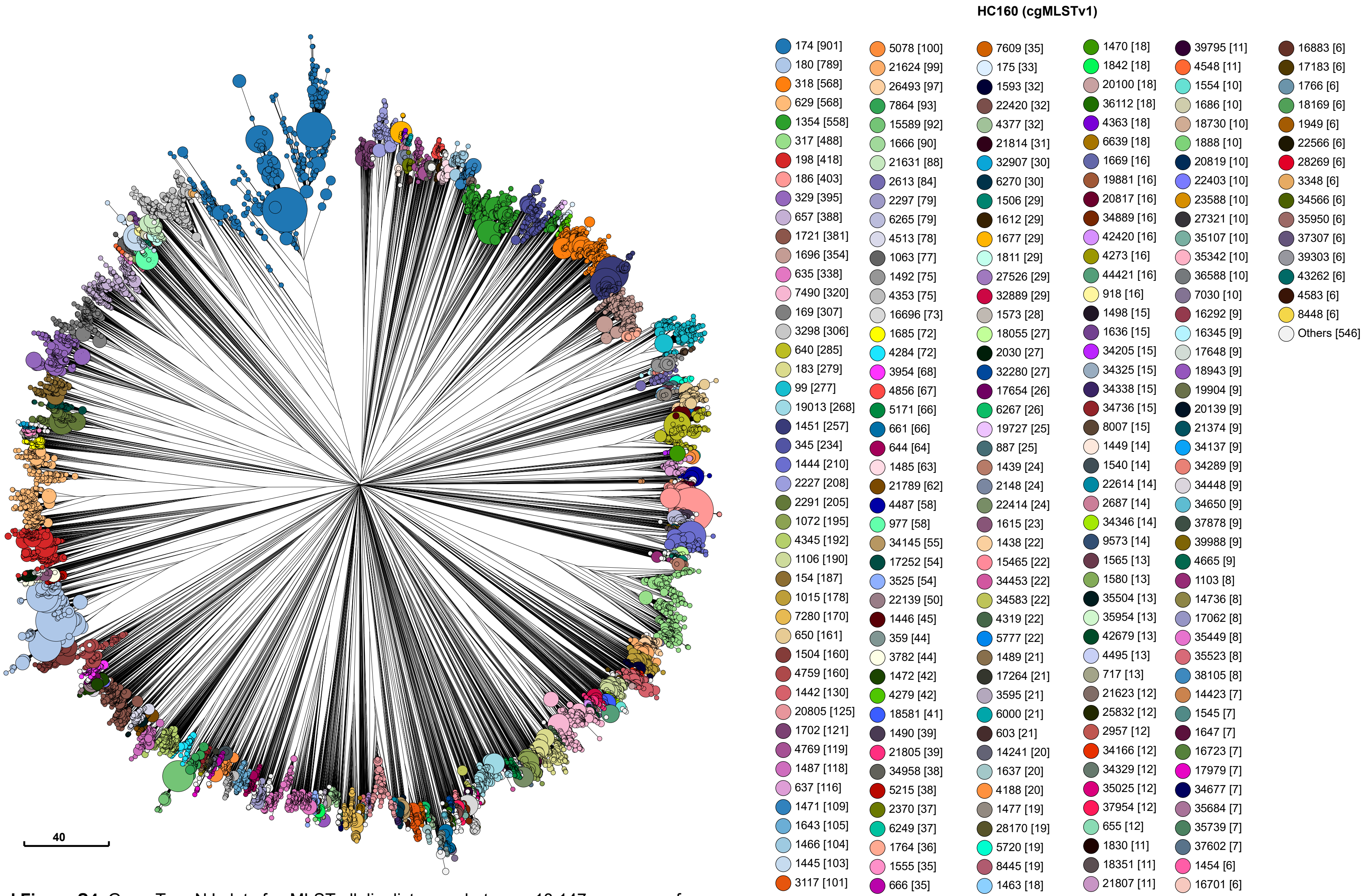


**Supplemental Figure S4.** GrapeTree NJ plot of cgMLST allelic distances between 18,147 genomes of *Streptococcus pneumonia* which had been assigned to GSPC clusters [107]. An interactive version of this figure may be examined at https://enterobase.warwick.ac.uk/ms_tree?tree_id=67948 and color-coded by HC100 cluster level, legacy Clonal Complex (CC), GPSC designation or by other metadata stored for those genomes within EnteroBase. Key Legend: HC160 cluster numbers and their color-codes. Legend: Measure of 40 allelic distances.
