## Supplementary material for "EnteroBase: Hierarchical clustering of 100,000s of bacterial genomes into species/sub-species and populations": Figure S5

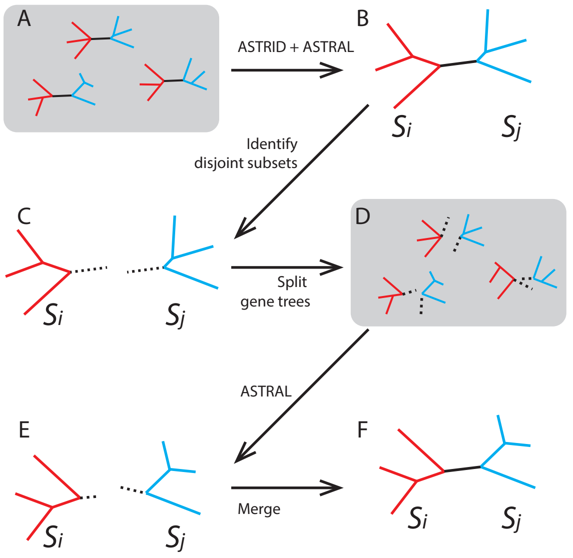


**Supplemental Figure S5.** Cartoon of the workflow of cgMLSA (https://github.com/zheminzhou/cgMLSA). (A) Construct gene trees for each core gene with RaXML. (B) Build an initial guide super-tree with ASTRID and calculate local branch supports with ASTRAL. (C) Split the initial guide tree into pairs of disjoint subsets at each highly supported branch, and append an artificial branch (dotted black line) to both members of the subset to allow later reconstruction of this branch. (D) Split all gene trees into the same sets of genomes for each pair of disjoint subsets. (E) Calculate a constraint super-tree for each subset with ASTRAL. (F) Construct the final species tree by merging constraint trees via the connections marked in C.
