## Supplementary material for "EnteroBase: Hierarchical clustering of 100,000s of bacterial genomes into species/sub-species and populations": Table S4

**Supplemental Table S4.** Taxonomic conflicts in *Vibrio*

| Species | HC1090 cluster | ANI95% (# genomes) | Problem strain(s) | Problem | Type Strains ignored |
|---|---|---|---|---|---|
| *V. cholerae* | 1 | 0 | *V. albensis* | Wrong taxonomy | *V. albensis* |
| *V. anguillarum* | 24 | 308 | *V.ordalii*<br>*V. qinghaiensis* | Wrong taxonomy | *V. ordalii*<br>*V. qinghaiensis* |
| *V. mediterranei* | 39 | 321 | *V. shilonii*<br>*V. barjaei* | Wrong taxonomy | *V. shilonii*<br>*V. barjaei* |
| *V. alginolyticus* | 69 | 41 (139)<br>106 (49) | ANI_106 | ANI_106 contains 10 *diabolicus*, 2 *antiquarius*, 1 *chemaguriensis,* 2 *parahaemolyticus*, 1 *cholerae,* 15 others | *V. antiquarius*<br>*V. chemaguriensis*<br>*V. diabolicus* |
| *V. splendidus* | 166 | 112 (97), 4049(6) | Two ANI clusters | | |
| *V. jasicida* | 180 | 393 (20), 3991 (5) | *V. inhibens*<br>*V. hyugaensis* | ANI_3991 is *V. hyugaensis.* ANI_393 includes 1 *V. inhibens*. | *V. inhibens*<br>*V. hyugaensis* |
| *V. owensii* | 1301 | 394 (38) | *V. communis* | Wrong taxonomy | *V. communis* |
| *V.* HC1090_1339 | 1339 | 309 (13)<br>2114 (2) | V. *atlanticus*<br>V. *tasmaniensis* | Mixed taxonomy | V. *atlanticus*<br>V. *tasmaniensis* |
| *V. chagasii* | 1478, 6298 | 86 (14)<br>2281 (2) | Two contradictory pairs of HierCC and ANI clusters | | |
| *V. tubiashii* | 1512 | 657 (2)<br>2278 (1) | Two ANI clusters | | |
| *V.* HC1090_1711 | 1711 | 2843 (1)<br>4775 (2) | New taxon | | |
| *V.* HC1090_2595<br>*V. hepatarius* | 2595<br>5006 | 3020 (1)<br>4588 (1) | Two *V. hepatarius* strains in related HC1090/ANI clusters | | |
| *V. crassostreae* | 2609 | 113 (44)<br>389 (2)<br>2123 (56)<br>3339 (1)<br>4428 (1) | Five ANI clusters | *V. celticus* & *V. coralliirubri* type strains in ANI_113 | *V. celticus*<br>*V. coralliirubri* |
| *V.* HC1090_2627 | 2627 | 320 (20) | *V. kanaloae*<br>*V. toranzoniae* | Two species and their type strains in one HC1090/ANI cluster | *V. kanaloae*<br>*V. toranzoniae* |
| *V. lentus* | 3503 | 1271 (1)<br>3729 (57)<br>4120 (1) | Three ANI clusters | No type strain | |
| *V. maritimis* | 3827 | 3903 (1)<br>4774 (1) | Two ANI clusters | | |
| *V. gazogenes*<br>*V.* HC1090_7553 | 4486<br>7553 | 274 (1)<br>2108 (1) | 2 *V. gazogenes* type strains in related HC1090/ANI clusters | | |
| *V. mexicanus* | 5181 | 644 (1)<br>2297 (1) | Two ANI clusters | | |
